# JNK pathway suppression drives resistance to combination endocrine therapy and CDK4/6 inhibition in ER+ breast cancer

**DOI:** 10.1101/2025.01.08.631992

**Authors:** Sarah Alexandrou, Christine S. Lee, Kristine J. Fernandez, Celine E. Wiharja, Leila Eshraghi, John Reeves, Daniel A. Reed, Neil Portman, Zoe Phan, Heloisa H. Milioli, Iva Nikolic, Antonia L. Cadell, David R. Croucher, Kaylene J. Simpson, Elgene Lim, Theresa E. Hickey, Ewan K. A. Millar, Carla L. Alves, Henrik J. Ditzel, C. Elizabeth Caldon

## Abstract

**Purpose:** Endocrine therapy in combination with CDK4/6 inhibition doubles the progression-free survival of patients with advanced ER+ breast cancer, but resistance is inevitable, leaving patients with limited treatment options.

**Experimental Design:** We performed unbiased genome-wide CRISPR/Cas9 knockout screens using ER+ breast cancer cells to identify novel drivers of resistance to combination endocrine therapy (tamoxifen) and CDK4/6 inhibitor (palbociclib) treatment. Screen hits were validated by CRISPR/Cas9 knockout models, mechanistic analyses and evaluation of patient samples.

**Results:** Our screens identified the inactivation of JNK signalling, including loss of the kinase *MAP2K7*, as a key driver of combination resistance. We developed multiple CRISPR/Cas9 knockout ER+ breast cancer cell lines (MCF-7 and T-47D) to investigate the effects of *MAP2K7, MAPK8* and *MAPK9* loss. *MAP2K7* knockout increased metastatic burden *in vivo* and led to impaired JNK-mediated stress responses, as well as promoting cell survival and reducing senescence entry following endocrine therapy and CDK4/6 inhibitor treatment. Mechanistically, this occurred via loss of the AP-1 transcription factor c-JUN, leading to an attenuated response to combination endocrine therapy plus CDK4/6 inhibition.

Furthermore, we analysed ER+ advanced breast cancer patient cohorts and found that inactivation of the JNK pathway was associated with increased metastatic burden, and low pJNK^T183/Y185^ activity correlated with a poorer response to systemic endocrine and CDK4/6 inhibitor therapies.

**Conclusions:** Overall, we demonstrate that suppression of JNK signalling enables persistent growth during combined endocrine therapy and CDK4/6 inhibition. Furthermore, our data provide a pre-clinical rationale to screen patients’ tumours for JNK signalling deficiency prior to receiving combined endocrine therapy and CDK4/6 inhibition.

**STATEMENT OF TRANSLATIONAL RELEVANCE:** Resistance to CDK4/6 inhibitors in the context of endocrine therapy resistance presents an urgent clinical challenge for the management of estrogen receptor positive (ER+) breast cancer. However, the mechanisms driving resistance to this therapeutic combination remain poorly understood. Here, we have identified inactivation of JNK signalling, specifically loss of the JNK kinase *MAP2K7*, as a major determinant of resistance to combined endocrine therapy and CDK4/6 inhibition in ER+ breast cancer. We uncovered that *MAP2K7* loss augments tumour survival *in vitro* and *in vivo.* This occurs via disruption of activator protein-1 (AP-1) transcription factors, and thus prevents the induction of therapy-induced senescence and JNK-induced stress response. These findings reveal a critical tumour suppressor role for the JNK pathway in ER+ breast cancer, highlighting the importance of identifying patients with deficient JNK signalling and cautioning against the development of JNK inhibitors for this setting.

## INTRODUCTION

Approximately 75% of all breast cancers express the estrogen receptor (ER), and patients commonly receive adjuvant endocrine therapy to target the ER (tamoxifen, aromatase inhibitors, fulvestrant). However, ∼35% of these patients eventually develop endocrine-resistant disease. Once metastatic, endocrine resistant ER+ breast cancer becomes incurable and accounts for ∼50% of breast cancer deaths [1]. The recent addition of cyclin-dependent kinase 4/6 (CDK4/6) inhibitors (e.g. palbociclib, ribociclib, abemaciclib) to endocrine therapy offers patients with resistant disease a new line of treatment. Combination therapy has doubled the progression-free survival of patients [2–4] and it is expected that metastatic ER+ breast cancer patients in countries with CDK4/6 inhibitor approval will likely receive combination endocrine therapy plus CDK4/6 inhibition during the course of their treatment. While highly effective, this combination therapy is not curative, and disease inevitably progresses [5].

Resistance to dual endocrine-CDK4/6 inhibitor therapy has emerged with the increased use of this combination. This represents the next major clinical challenge in ER+ breast cancer as there are currently limited therapeutic strategies upon CDK4/6 inhibitor failure. Contemporary studies have identified putative “drivers” for CDK4/6 inhibitor resistance such as Rb loss, FAT1 loss, CDK6 gain or cyclin E1 gain [6–8]. However, these mechanisms have been validated in only a small number of patients, possibly due to the predominance of pre-clinical studies using CDK4/6 inhibitors as single agents. Although pre-clinical studies of endocrine monotherapy resistance and CDK4/6 inhibitor monotherapy resistance are available, pre-clinical studies of combination endocrine-CDK4/6 inhibitor resistance mechanisms are rare. Therefore, there is an urgent need for studies utilising combination therapies that mimic clinical practice to be employed to study resistance mechanisms.

In this study we identified genes whose loss-of-function leads to insensitivity to combination endocrine and CDK4/6 inhibitor therapy in MCF-7 ER+ breast cancer cells using human genome-wide CRISPR/Cas9 screens. We identified the c-Jun N-terminal kinase (JNK) pathway as a major determinant of resistance to endocrine therapy and CDK4/6 inhibition in ER+ breast cancer. The lead hit was *MAP2K7*, a kinase that directly phosphorylates JNK to control JNK pathway activation. *In vitro* validation using guide RNAs (gRNAs) confirmed that loss of the JNK pathway, particularly *MAP2K7*, impaired the anti-proliferative and pro-senescent response of cells to endocrine therapy and CDK4/6 inhibition. Moreover, these cells have an impaired JNK-mediated stress response. Finally, we demonstrated that low expression of *MAP2K7* and pJNK^T183/Y185^ was associated with poor outcomes in metastatic ER+ breast cancer patients treated with endocrine therapy and CDK4/6 inhibitors. Our findings strongly suggest that the JNK pathway acts as a tumour suppressor in ER+ breast cancer, and cautions against the use of JNK inhibitors in this setting.

## MATERIALS AND METHODS

### Cell lines and drug treatment

MCF-7 and T-47D cells were cultured in phenol-red free RPMI 1640 (Gibco) supplemented with 5% charcoal-stripped fetal bovine serum (FBS), 1% penicillin/streptomycin (Invitrogen) and 10 pM 17β-estradiol (Sigma-Aldrich; resuspended in absolute ethanol (EtOH)). HEK293T cells were cultured in DMEM (Gibco) supplemented with 10% FBS. All cell lines were cultured for less than 6 months after short-tandem repeat profiling (Garvan Molecular Genetics, Garvan Institute of Medical Research (GIMR)), and routinely tested negative for mycoplasma contamination. Cells were cultured under 5% CO_2_ in a humidified incubator at 37°C.

The MCF-7 and T-47D *MAP2K7*^-/-^, MCF-7 *MAPK8*^-/-^ and MCF-7 *MAPK9*^-/-^ cell lines were generated from MCF-7-Cas9- or T-47D-Cas9-expressing cells, and transduced with gRNAs targeting MKK7 (*MAP2K7*), JNK1 (*MAPK8*) and JNK2 (*MAPK9*). *MAP2K7, MAPK8* and *MAPK9* gRNAs in pLentiGuide-Puro were purchased from GenScript. The gRNA sequences were: MKK7_sg1 ACGGGCTACCTGACCATCGG, MKK7_sg3 CATTCTGGGCAAGATGACAG, MAPK8 TCGCTACTACAGAGCACCCG and MAPK9 AATGGATGCTAACTTATGTC. Sequences for the *MAP2K7* gRNAs were chosen based on high enrichment in the pooled CRISPR/Cas9 screen.

Cells were treated with the following: tamoxifen (Sigma-Aldrich; resuspended in tetrahydrofuran), fulvestrant (Sigma-Aldrich; resuspended in EtOH), palbociclib (Selleckchem; resuspended in water), anisomycin (Sigma-Aldrich; resuspended in dimethyl sulfoxide), and puromycin (Sigma-Aldrich; resuspended in water).

### *In vivo* analysis of metastatic burden

#### Xenograft model

All animal work was performed in compliance with the Australian Code of Practice for the Care and Use of Animals for Scientific Purposes (National Health and Medical Research Council). The protocol and study end points were approved by the St. Vincent’s Health Precinct and GIMR Animal Ethics Committee (ARA 21/09). Immunocompromised NOD-SCID-IL2γ^-/-^5-6 week old female mice were housed in ventilated cages and specific pathogen-free conditions in a 12:12 hour light:dark cycle with food and water given *ad libitum*. After an acclimatisation period of a minimum of seven days, 10 mice per arm were injected via the tail vein with 1.5 x 10^6^ MCF-7 pLenti and *MAP2K7*^-/-^ (MKK7_sg3) cells. Cell growth was supported by 17β-estradiol (2.5 µg/mL), administered in the drinking water. Mice were monitored for 9 weeks, with twice weekly bladder palpation to monitor for estradiol toxicity, until ethical endpoint for metastatic burden was reached. Mice were euthanised with isoflurane anaesthetisation and cervical dislocation, and the lungs were harvested and fixed for 24 hours in 10% neutral buffered formalin before being transferred to 70% ethanol.

#### Immunohistochemistry (IHC) and quantification (Cytokeratin)

Formalin-fixed paraffin-embedded (FFPE) sections from mouse lung tissue were cut with a microtome, mounted on Epredia SuperFrost slides and dried at 60°C. Sections were deparaffinised and stained using the Bond RX Automated Stainer (Leica Biosystems). For Cytokeratin staining, heat-induced antigen retrieval was performed at pH9 (Bond Epitope Retrieval solution 2, Leica Biosystems), 100°C for 30 minutes, before antibody incubation (1:500; MA1-12594; Thermo Fisher Scientific) for 30 minutes. Detection was performed with diaminobenzidine (DAB, EnVision Detection Systems, Agilent Dako) and slides were counterstained with haematoxylin. Slides were imaged using a slide scanner (AperioCS2, Leica Biosystems), and images were analysed using QuPath software (version 0.4.4) by manually selecting positively stained regions [9].

### Lentiviral production and generation of stable *MAP2K7^-/-^, MAPK8^-/-^ and MAPK9^-/-^* cell lines

HEK293T cells were used to package virus. Plasmids were extracted using the PureLink HiPure Plasmid Filter Maxiprep Kit (Invitrogen) as per the manufacturer’s instructions. 4 x 10^6^ HEK293T cells were seeded into 10 cm^2^ plates and transfected with either *MAP2K7*, *MAPK8,* or *MAPK9* gRNA plasmids, along with packaging plasmids pMDLg/pRRE, pRSV-REV, and pMD2.G. Lentiviral-containing medium was collected 48 hours after transfection and filtered using a 0.45 µm filter. MCF-7-Cas9 or T-47D-Cas9 cells were then immediately infected with lentivirus at a 1:2 dilution with 8 µg/mL polybrene (Sigma-Aldrich). Selection with 2 µg/mL puromycin occurred 48 hours after transduction to generate stably expressing MCF-7-Cas9 and T47-D-Cas9 cell lines, including pLenti (empty vector), MKK7_sg1 and MKK7_sg3, and MCF-7-Cas9 *MAPK8*^-/-^ and *MAPK9*^-/-^ cell lines. Cells were then cultured with a maintenance dose of 1 µg/mL puromycin.

### Cell proliferation, clonogenic assays and synchronisation treatments

Metabolic assays were performed in 96-well plates with cells seeded at 1 x 10^3^ per well. Cells were treated with drug or vehicle for up to 5 days at the concentrations indicated in the figure legends, and metabolic rate was assessed using AlamarBlue (Thermo Fisher Scientific). Half-maximal inhibitory concentration (IC_50_) values were determined using GraphPad Prism software (version 10). Live-cell imaging of cell number was performed using the Incucyte S3 (Live-Cell imaging and Analysis System; Sartorius). 2 x 10^3^ cells per well were seeded in 96-well plates and imaged over 7 days, with 2 fields of view per well. Images were analysed using IncuCyte ZOOM Software (version 2020C).

For 90-day growth-rate assays, MCF-7 cells were seeded at 4.8 x 10^6^ in T150 flasks. Following a 24-hour incubation to allow cells to attach, cells were continuously treated with 500 nM palbociclib or 500 nM tamoxifen plus 250 nM palbociclib. Media and drugs were changed every 3-4 days and cells were passaged at 80% confluency. Live cell counts using trypan blue staining (Gibco) were used to calculate cell number to determine growth rates.

Clonogenic assays were performed in 12-well plates with cells seeded at 3 x 10^3^ cells per well. Drugs were replenished every 3-4 days, for up to 3 weeks. Colonies were fixed with trichloroacetic acid (16%) and stained with 0.1% – 0.5% crystal violet (Sigma-Aldrich). Plates were scanned using Epson Perfection V800 photo scanner at 1200 dots per inch. Colony area was quantitated using ImageJ software (version 2.1.0/1.53c).

MCF-7 and T-47D pLenti and *MAP2K7*^-/-^ cells synchronized at G_1_/S phase with 48 hours 10 nM fulvestrant were released into medium supplemented with 100 nM 17β-estradiol for 12 and 24 hours.

### Immunoblotting

Cells were lysed in ice-cold Normal Lysis Buffer (10% (v/v) glycerol, 1.2% (w/v) HEPES, 1% (w/v) sodium acid pyrophosphatase, 1% (v/v) Triton X-100, 0.8% (w/v) sodium chloride, 0.4% (w/v) sodium fluoride, 0.04% EGTA, 0.03% (w/v) magnesium chloride) supplemented with 200 µM sodium orthovanadate, 1 mM dithiothreitol, 50 µL/mL protease inhibitor cocktail (Sigma-Aldrich) and 10 µg/mL MG132. Cell lysates were separated on 4-12% Bis-Tris gels (Invitrogen) as previously described [10].

Primary antibodies used at a 1:1000 dilution were MKK7 (SDC20-87; Invitrogen), MKK4 (PA5-96776; Invitrogen), phospho-JNK^T183/Y185^ (9251; Cell Signalling Technology), JNK (9252; Cell Signalling Technology), JNK1 (3708; Cell Signalling Technology), ERα (8644; Cell Signalling Technology), phospho-cJUN^Ser63^ (sc-822; Santa Cruz Biotechnology), cJUN (9165; Cell Signalling Technology) and JUND (5000; Cell Signalling Technology). GAPDH was used at 1:15000 dilution (6C5; Santa Cruz Biotechnology). Secondary antibodies, chemiluminescence and densitometry were performed as previously described [10].

### Quantitative real-time polymerase chain reaction (PCR) analysis

Total RNA was extracted from MCF-7 and T-47D pLenti and *MAP2K7*^-/-^ cell pellets using RNeasy Plus Mini kit (Qiagen). Reverse transcription was performed using the High-Capacity cDNA Reverse Transcription Kit (Applied Biosystems) to detect mRNA. *MAP2K7* (hs00178198_m1), *JUN*(hs01103582_s1), *JUND* (hs04187679_s1), *ATF3* (hs00231069_m1) and *RPLP0* (hs_00420895_gh) Taqman probes (Thermo Fisher Scientific) were used to analyse mRNA expression levels using a QuantStudio7 Flex Real-Time PCR System. Relative gene expression was assessed using the ΔΔCycle threshold method [11], where expression was normalised to *RPLP0*.

### Senescence associated **β**-galactosidase assay

Senescence was assessed by visualising β-galactosidase activity using the Senescence β- Galactosidase Staining Kit as per the manufacturer’s instructions (Cell Signalling Technology). β-galactosidase activity was imaged using the Leica DFC295 microscope, and the proportion of positive and negative β-galactosidase-stained cells across 3-8 fields of view were quantified using ImageJ Cell Counter.

### Flow cytometry

Ethanol-fixed cells were stained with 1 µg/mL propidium iodide (PI; Sigma-Aldrich) overnight, and incubated with 0.5 mg/mL RNaseA (Sigma-Aldrich) for 2-5 hours. Cells were then analysed on a BD FACSCanto II (BD Biosciences). DNA histograms contained ∼30,000 events, and cell cycle distribution was analysed using FlowJo (version 10.6.1) and ModFit (Verity Software House, version 6.0).

Apoptosis was assessed using the Annexin V-FITC Apoptosis Kit (#K101; BioVision) on the BD FACSCanto II. Cell death by apoptosis was measured by quantifying the proportion of FITC positive/PI negative (early apoptosis) and FITC positive/PI positive (late apoptosis) cells on >30,000 events using FlowJo.

### Genome-wide pooled CRISPR/Cas9 screen

#### Cas9 cell line generation

MCF-7 and T-47D cells stably expressing Cas9 were generated by transducing cells with Cas9-mCherry lentivirus (Addgene #70182; obtained from the Victorian Centre for Functional Genomics (VCFG)) at a multiplicity of infection (MOI) of 0.3. Cas9-mCherry cells were FACS sorted on a BD FACSAria III (BD Biosciences) for the population with the top 10% highest mCherry expression.

#### Positive selection pooled knockout screen

The human genome-wide pooled Brunello library (containing 77,441 sgRNAs) was provided by the VCFG (Addgene #73178). MCF-7-Cas9 cells were infected with the Brunello library at an MOI of 0.3 using 8 µg/mL polybrene. In parallel, MCF-7-Cas9 cells were used as mock controls and treated only with 8 µg/mL polybrene. 1 x 10^6^ cells/well were plated in 12-well plates and spin-infected (30 minutes, 30°C, 1,200 × g) to increase lentiviral transduction efficiency. 72 hours after infection, Brunello-library-transduced and mock-transduced cells were selected with 2 µg/mL puromycin. Functional titre was optimised for the Brunello library to achieve a 22% transduction efficiency.

Following selection, the T_0_ baseline timepoint was collected at 1,000-fold representation (7.6 x 10^7^ cells). To ensure the complexity of the Brunello library was maintained at 500-fold representation during treatment (aiming for 500 copies per sgRNA), 3.8 x 10^7^ transduced cells per treatment group was needed at the start of the experiment. Following a 48 hour incubation to allow cells to attach, cells were continuously treated with 500 nM tamoxifen plus 250 nM palbociclib (early: 6 week and late: 10 well timepoint) or 500 nM palbociclib alone (early: 2 week and late: 4 week timepoint), supplemented with 1 µg/mL puromycin. Media and drugs were changed every 3-4 days and cells were passaged at 80% confluency. Importantly, no cells were discarded they were continuously passaged. Pellets of cells were collected for genomic DNA (gDNA) isolation and library preparation at early and late timepoints for each treatment arm.

#### CRISPR screen sample preparation and sgRNA library sequencing

gDNA was extracted from cell pellets of the early and late timepoints using Midi and Maxi Qiagen Blood and Cell Culture Kits (Qiagen) as per the manufacturer’s instructions. PCR amplification and purification of DNA using AMPure beads were then performed as per instructions from VCFG. Before sequencing, Qubit DNA assay was performed for quantification, and a small amount of each sample was run on a 2% agarose gel to confirm the 345 kb PCR product. A minimum of 500 ng PCR product per sample (or ∼50 ng specific PCR product) was required for a target of 40 million reads.

Samples were sequenced using 75 base pair single-end sequencing at the Molecular Genomics Core facility (Peter MacCallum Cancer Centre) using the Illumina NextSeq 500 platform. Raw FASTQ files were analysed using MAGeCK-VISPR [12] and mapped to the human genome-wide pooled Brunello library (available from https://sourceforge.net/p/mageck/wiki/libraries/). The number of mapped reads per sample ranged from 13 million to 19 million, with an average map-ability of 66% (Supplementary Figure 1A). Sequencing revealed clear differences in the representation of sgRNAs as measured by the Gini index, which is a determinant of inequality among samples (Supplementary Figure 1B). MAGeCK-MLE (version 0.5.9.3) was used to rank and sort sgRNAs by false discovery rate (FDR) <0.5 and the results were visualised in VISPR (version 0.4.15) (Supplementary Table 1). Gene set enrichment analysis (GSEA) was performed with MSigDB using Canonical Pathways (Supplementary Table 2).

### RNAseq data analysis, normalisation and differentially expressed gene identification

MCF-7 and T-47D pLenti and *MAP2K7^-^*^/-^ (MKK7_sg3) cells lines were treated with either tamoxifen or fulvestrant plus palbociclib for 48 hours. Total RNA was extracted from cell pellets as described above. 1 μg RNA was used to prepare libraries using the KAPA mRNA HyperPrep Kit (Roche). Reads were sequenced on a NovaSeq 6000 instrument (Illumina), using 150 base pair paired-end sequencing chemistry, by the Sequencing Platform (GIMR). FASTQ files were quality checked by ‘FastQC’ (version 0.11.9) [13]. Sequence adaptors and quality filtering was performed using ‘*AdapterRemoval*’ (version 2.3.1) and processed paired-end FASTQ reads were aligned to the reference genome assembly ‘GRCh38_2020- A_build’ [14] using STAR (version 2.7.8) [15] with default parameters. Gene expression was quantified by counting the number of reads aligned to each Ensembl gene model using ‘featureCounts’ (version 2.0.1) [16] available through the package RSubread (https://bioconductor.org/packages/release/bioc/html/Rsubread.html).

Differentially expressed genes (DEGs) between the different treatment groups were identified using the *limma-voom* method (version 3.46.0) [17, 18]. To normalise read counts according to library size differences between the samples, the Trimmed Mean of M-values normalisation method from *edgeR* (version 3.32.1) was applied [19]. To identify whether the RNAseq library preparation date contributed to variation in gene expression patterns, multidimensional scaling (MDS) was performed to visualise the summary of gene expression for all libraries. Genes with low expression were filtered out prior to DEG analysis and only genes with at least 1 count per million reads in the three replicate samples were kept. Genes with FDR adjusted p-value[<0.05 were considered DEGs (Supplementary Table 3). Genes that were upregulated or downregulated more than 1-fold with each drug treatment (tamoxifen or fulvestrant plus palbociclib) in pLenti cells were identified. These lists were then screened for those genes which showed >0.25 fold less regulation in *MAP2K7*^-/-^ cells than in pLenti cells. Genes that showed reduced upregulation or downregulation in *MAP2K7*^-/-^ cells were analysed by GSEA against the Hallmark datasets using ShinyGo [20]. Data was visualised using Excel.

Transcription factors [21] that had altered expression in *MAP2K7*^-/-^ cells were identified using a combined cut-off of log fold change >0.5 between pLenti and *MAP2K7*^-/-^ cells. Venn network plots were prepared with EVenn [22].

### Clinical samples

#### Patient demographics and tumour samples

A cohort of 101 metastatic ER+ breast cancer patients treated with combined endocrine therapy and CDK4/6 inhibition with clinical follow-up data were selected as previously described [23]. Of these, 81 had available FFPE material from metastatic lesions for IHC analysis, following which 3 were excluded for lack of identifiable tumour tissue in the sample (Supplementary Figure 6A).

#### Phospho-JNK^T183/Y185^ IHC and quantification

FFPE sections from metastatic biopsies were cut with a microtome, mounted on ChemMateTM Capillary Gap Slides (Dako), dried at 60°C, deparaffinised, and hydrated. Sections were stained using the Bond RX Automated Stainer (Leica Biosystems). For pJNK^T183/Y185^ staining, heat-induced antigen retrieval was performed at pH6 (Bond Epitope Retrieval solution 1, Leica Biosystems), 100°C for 20 minutes, before antibody incubation (1:200; AF1205; R&D Systems) for 60 minutes. Detection was performed with DAB (Bond Polymer Refine Detection, Leica Biosystems) and slides were counterstained with haematoxylin. Slides were imaged using the AperioaCS2 slide scanner, and images were analysed using QuPath [9]. A semi-quantitative H-score was obtained by the sum of the percentage of the tumour cells (0-100%) for each staining intensity (0-3), giving a range of 0 to 300 [24]. H-scores were binned to identify the distribution of expression. Based on the overall distribution of pJNK^T183/Y185^ activity in the cohort (n=78), thresholds were selected to approximate an even distribution across the three groups: low expression (H-score <195, n=25), medium expression (H-score >195 and <246, n=26), and high expression (H-score >246, n=27) (Supplementary Figure 6B).

### In silico datasets

#### Primary and metastatic ER+ breast cancer datasets

Gene expression data was downloaded from The Cancer Genome Atlas (TCGA) [25, 26] through cBioPortal [27, 28] in January and December 2024. The following breast cancer patient cohorts were analysed: Molecular Taxonomy of Breast Cancer International Consortium (METABRIC) [29], TCGA Firehose Legacy [30], and Metastatic Breast Cancer Project (MBCP; provisional dataset) [31]. Each dataset was analysed independently, and ER status was defined as positive by IHC (TCGA and METABRIC) and ‘pathology reports’ (MBCP). JNK1, JNK2, and pJNK protein expression from Reverse-Phase Protein Arrays and Clinical Proteomic Tumour Analysis Consortium [32] data were also obtained from TCGA and UALCAN [33].

#### Tamoxifen-treated ER+ breast cancers

Array-based gene expression and clinical data from a cohort of tamoxifen-treated ER+ patients (GSE9893) was downloaded from https://www.ncbi.nlm.nih.gov/geo/query/acc.cgi?acc=GSE9893 [34]. mRNA expression of Jetset probes [35] was stratified based on metastatic burden. Comparisons were performed using Mann-Whitney unpaired t-test.

#### Neoadjuvant palbociclib treatment

Gene expression arrays and clinical data from a cohort of early-stage ER+ breast cancer patients treated with pre-operative palbociclib for 2 weeks (n=72 samples) were accessed from http://microarrays.curie.fr/publications/U981-GustaveRoussy/pop/ [36]. mRNA expression was stratified into responders and non-responders based on anti-proliferative effect, with responders defined as those showing a significant reduction in proliferation markers.

### Statistical analysis

Statistical analysis was performed using GraphPad Prism (version 10). Pairs of datasets were compared using unpaired two-tailed t-tests. Groups of data were compared using one-way ANOVA followed by multiple comparisons with Tukey’s test, two-way ANOVA followed by multiple comparisons with Tukey’s test or mixed effects analysis followed by multiple comparisons with Tukey’s test. Pearson’s analysis was performed to examine correlation between datasets. Kaplan-Meier survival analysis was performed with the Log-rank (Mantel-Cox) test, and hazard ratios were calculated. Violin plots are presented from minimum to maximum, with the range extending from 1^st^ to 3^rd^ quartile, and the line in the middle indicating the median. All experiments were performed in biological triplicate unless otherwise specified. Data are presented as mean ± standard error of the mean (SEM).

## RESULTS

### Genome-wide CRISPR/Cas9 screen identifies JNK pathway suppression as a driver of endocrine-CDK4/6 inhibitor resistance

To identify novel genes conferring resistance to combination endocrine therapy and CDK4/6 inhibition in ER+ breast cancer, we performed positive selection (enrichment) genome-wide knockout CRISPR/Cas9 screens in ER+ MCF7 breast cancer cells (n=2 biological replicates; Figure 1A). MCF-7 cells engineered to stably express Cas9 were transduced with the Brunello lentiviral pooled library, which targets 19,114 genes [37]. Following puromycin selection, transduced cells were treated with the combination of 500 nM tamoxifen and 250 nM palbociclib for 6 (early) and 12 (late) weeks, or 500 nM palbociclib monotherapy for 2 (early) and 4 (late) weeks (approximate IC_50_ doses; Supplementary Figure 1C, D). The combination of tamoxifen and palbociclib was chosen as CDK4/6 inhibition with tamoxifen is currently used clinically and has reported improved survival benefit in patients compared to endocrine therapy alone [38]. Moreover, treatment refractory cells emerge within 12 weeks on this treatment combination (Supplementary Figure 1E), yielding sufficient material for downstream analysis.

**Figure 1:**
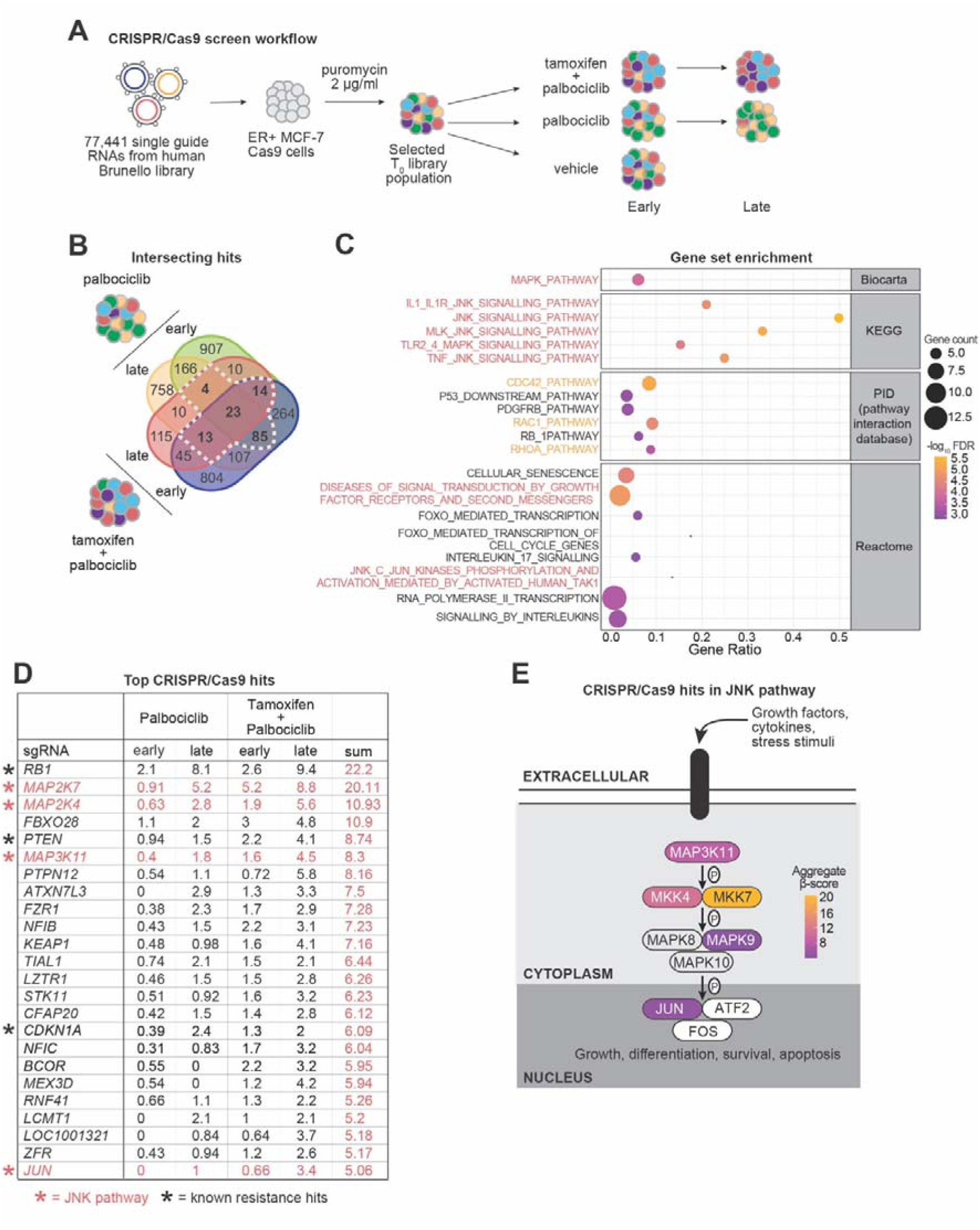
CRISPR/Cas9 screen identifies JNK pathway deficiency in endocrine-CDK4/6 inhibitor resistance. **(A)** A positive selection CRISPR/Cas9 screen was conducted in ER+ MCF-7-Cas9 expressing cells using the human Brunello library. Cells were treated with the combination of 500 nM tamoxifen and 250 nM palbociclib (early: 6 week and late: 10 week timepoint), or 500 nM monotherapy palbociclib (early: 2 week and late: 4 week timepoint) or vehicle (tetrahydrofuran) before genomic DNA was collected and sequenced. **(B)** Venn diagram showing single guide RNAs (sgRNAs) that increased following treatment with palbociclib and tamoxifen + palbociclib. sgRNAs selected with a false discovery rate <0.5, and occurrence in ≥3 screens, where at least one screen was with combination therapy. Selected sgRNAs indicated with dashed white line. **(C)** Gene set enrichment analysis of commonly downregulated genes with ≥2.5 aggregate β-score. **(D)** Top sgRNAs enriched following treatment with palbociclib and tamoxifen + palbociclib, and corresponding β-scores (sgRNAs with aggregate β-score ≥5 are shown). Orange * indicates sgRNAs in the JNK pathway. Black * indicates sgRNAs described in the literature to drive CDK4/6 inhibitor resistance. **(E)** Schematic of the JNK signalling pathway and enriched sgRNAs, as well as aggregate β- score scale. P indicates phosphorylation events.

Genomic DNA from each treatment condition was collected after 2/4 weeks (palbociclib arm) and 6/12 weeks (tamoxifen plus palbociclib arm), and sequenced to determine the sgRNA representation compared to the T_0_ sgRNA counts using MAGeCK-VISPR [12] (Supplementary Table 1). More than 1.2 x 10^7^ reads were detected per sample, with at least 60% mapped, and Gini analysis showed sgRNA enrichment with selective conditions (Supplementary Figure 1A, B). Candidate sgRNAs associated with resistance to tamoxifen and palbociclib were identified using the inclusion criteria of a FDR <0.5 and occurrence in at least three screens, including one with the combination therapy (Figure 1B).

We identified 139 genes and next interrogated this list for known and novel drivers of CDK4/6 inhibitor resistance. *RB1*, *CDKN1A* and *PTEN* were significantly enriched and have previously been described in the literature to confer resistance to CDK4/6 inhibitors [39–43] (Supplementary Table 2). GSEA was performed on the top genes (composite enrichment score ≥2.5), revealing the involvement of MAPK and JNK signalling pathways (Figure 1C). GTPases such as CDC42, Rac1 and Rhoa were also enriched (Figure 1C), and notably, JNK signalling is a key downstream component of these pathways [44, 45].

Overall, we observed a cluster of genes encoding proteins involved in JNK signalling. The JNK pathway involves a phosphorylation cascade that activates JNK, leading to activation of downstream pathways that modulate proliferation and cell survival. We observed significant depletion across the JNK cascade, including the upstream mediators MKK7 *(MAP2K7)*, MKK4 *(MAP2K4)*, and *MAP3K11*, the gene that encodes the JNK2 protein (*MAPK9*), and the downstream transcriptional mediator *JUN* (Figure 1D, E; Supplementary Table 2). In our CRISPR/Cas9 screen *RB1* was the most enriched gene, however the β-scores were very similar across the two treatment arms (palbociclib composite score: 10.2; tamoxifen plus palbociclib composite score: 12), indicating that *RB1* loss was primarily driven by CDK4/6 inhibition (Figure 1D). *MAP2K7* was identified as a top depleted target gene with an aggregate β-score of 20.11 (palbociclib composite score: 6.11; tamoxifen plus palbociclib composite score: 14) (Figure 1D, E). Interestingly, this result indicates that resistance to palbociclib alone or in combination with tamoxifen, is potentially due to *MAP2K7* loss and the disruption of JNK signalling. This was surprising as the role of the JNK pathway in cancer is frequently described as oncogenic, although it is reported to act as a tumour suppressor in certain cancer contexts [46].

### *MAP2K7* loss reduces pJNK^T183/Y185^ activity in ER+ breast cancer

Of the CRISPR/Cas9 target hits, *MAP2K7* was the most enriched sgRNA in the JNK pathway. *MAP2K4* was also highly enriched in our CRISPR screen however, it is known to activate p38 signalling in addition to JNK signalling [47]. This led us to hypothesise that *MAP2K7*/MKK7 is a novel and specific determinant of JNK signalling dysfunction in ER+ breast cancer.

To validate the effects of MKK7 loss in ER+ breast cancer, we engineered multiple *MAP2K7* CRISPR/Cas9 knockout cell lines in MCF-7 and T-47D cells (MKK7_1 and MKK7_3; Figure 2A, B). Knockout of *MAP2K7* resulted in >97% protein suppression in all *MAP2K7*^-/-^ MCF-7 and T-47D cell lines (Figure 2A, B), without altering MKK4 protein expression (Figure 2A, B; Supplementary Figure 2A-D).

**Figure 2:**
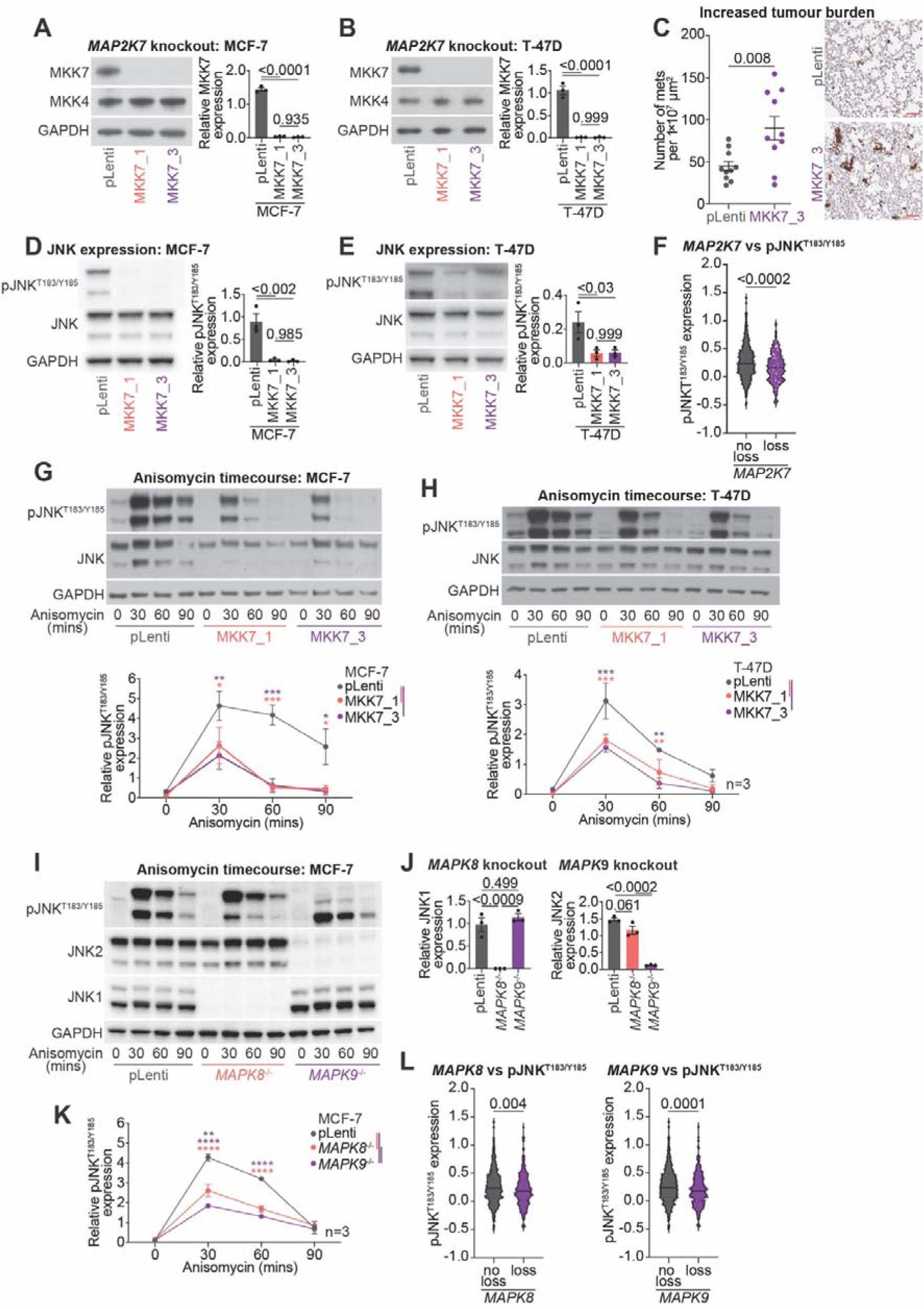
*MAP2K7* loss leads to reduced JNK phosphorylation and diminished induction of JNK-induced stress response. **(A)** Representative Western blot of MKK7, MKK4 and GAPDH in MCF-7 pLenti and *MAP2K7*^-/-^ (MKK7_1 and MKK7_3) cells (full length blots in Supplementary Figure 2B). Quantitation of MKK7 expression by densitometry. Band intensity was normalised to GAPDH. Data analysed by one-way ANOVA with Tukey’s multiple comparisons test. **(B)** Representative Western blot of MKK7, MKK4 and GAPDH in T-47D pLenti and *MAP2K7*^-/-^ (MKK7_1 and MKK7_3) cells (full length blots in Supplementary Figure 2D). Quantitation of MKK7 expression by densitometry. Band intensity was normalised to GAPDH. Data analysed by one-way ANOVA with Tukey’s multiple comparisons test. **(C)** Metastases to lung of MCF-7 pLenti and MKK7_3 xenografts (n=10 mice per arm) as measured by cytokeratin immunohistochemistry. Number of metastases quantitated per 1 x 10^7^ µm^2^ area and analysed by unpaired two-tailed t-test. Representative images shown, scale bar = 200 µm. **(D)** Representative Western blot of pJNK^T183/Y185^, JNK and GAPDH in MCF-7 pLenti and *MAP2K7*^-/-^ (MKK7_1 and MKK7_3) cells (full length blots in Supplementary Figure 2F). Quantitation of pJNK^T183/Y185^ activity by densitometry. Band intensity was normalised to GAPDH. Data analysed by one-way ANOVA with Tukey’s multiple comparisons test. **(E)** Representative Western blot of pJNK^T183/Y185^, JNK and GAPDH in T-47D pLenti and *MAP2K7*^-/-^ (MKK7_1 and MKK7_3) cells (full length blots in Supplementary Figure 2H). Quantitation of pJNK^T183/Y185^ activity by densitometry. Band intensity was normalised to GAPDH. Data analysed by one-way ANOVA with Tukey’s multiple comparisons test. **(F)** Expression of pJNK^T183/Y185^ in primary ER+ breast cancers from the TCGA cohort compared to *MAP2K7* copy number status. *MAP2K7* “no loss” includes cancers with amplified, gain or diploid status for *MAP2K7*. *MAP2K7* “loss” includes cancers with heterozygous or homozygous deletion of *MAP2K7*. Data analysed by unpaired two-tailed t-test. **(G)** Representative Western blot of pJNK^T183/Y185^, JNK and GAPDH in MCF-7 pLenti and *MAP2K7*^-/-^ (MKK7_1 and MKK7_3) cells treated with 300 nM anisomycin for 0, 30, 60, 90 minutes (full length blots in Supplementary Figure 2I). Quantitation of pJNK^T183/Y185^ activity by densitometry. Band intensity was normalised to GAPDH. Data analysed by two-way ANOVA with Tukey’s multiple comparisons test. **(H)** Representative Western blot of pJNK^T183/Y185^, JNK and GAPDH in T-47D pLenti and *MAP2K7*^-/-^ (MKK7_1 and MKK7_3) cells treated with 300 nM anisomycin for 0, 30, 60, 90 minutes (full length blots in Supplementary Figure 2J). Quantitation of pJNK^T183/Y185^ activity by densitometry. Band intensity was normalised to GAPDH. Data analysed by two-way ANOVA with mixed effects analysis and Tukey’s multiple comparisons test. **(I)** Representative Western blot of pJNK^T183/Y185^, JNK2, JNK1 and GAPDH in CRISPR/Cas9 MCF-7 cell lines: pLenti, *MAPK8*^-/-^ and *MAPK9*^-/-^. Cells were treated with 300 nM anisomycin for 0, 30, 60, 90 minutes (full length blots in Supplementary Figure 2K). **(J)** JNK1 and JNK2 expression was quantitated at 0 min timepoint. Band intensity was normalised to GAPDH. Data analysed by one-way ANOVA with Tukey’s multiple comparisons test. **(K)** Quantitation of pJNK^T183/Y185^ activity by densitometry. Band intensity was normalised to GAPDH. Data analysed by two-way ANOVA with Tukey’s multiple comparisons test. **(L)** Expression of pJNK^T183/Y185^ in primary ER+ breast cancers from the TCGA cohort compared to *MAPK8* or *MAPK9* copy number status. *MAPK8* or *MAPK9* “no loss” includes cancers with amplified, gain or diploid status for *MAPK8* or *MAPK9*. *MAPK8* or *MAPK9* “loss” includes cancers with heterozygous or homozygous deletion of *MAPK8* or *MAPK9*. Data analysed by unpaired two-tailed t-test.

Given that the JNK pathway can act as both a tumour suppressor and as an oncogene [48], we first assessed whether *MAP2K7* loss would increase tumour burden *in vivo*. We injected MCF-7 MKK7_3 knockout cells into the tail-vein of immunocompromised mice, with pLenti cells as a control. After 9 weeks, the lungs of mice were harvested and examined for micrometastases by IHC for human cytokeratin. Mice xenografted with MCF-7 *MAP2K7*^-/-^ cells incurred, on average, a 2-fold higher number of metastases than those xenografted with MCF-7 pLenti cells (p=0.008; Figure 2C), demonstrating that *MAP2K7* loss in these cells increased metastasis.

We next examined the impact of *MAP2K7* loss on JNK phosphorylation. Loss of *MAP2K7* resulted in a 75-97% reduction in endogenous pJNK^T183/Y185^ levels in MCF-7 and T-47D cells, indicating that MKK7 mediates a substantial proportion of JNK phosphorylation in proliferating ER+ breast cancer cells. Loss of *MAP2K7* did not alter total JNK protein expression in either MCF-7 or T-47D cells (Figure 2D, E; Supplementary Figure 2E-H). We confirmed the co-occurrence of *MAP2K7* deletion and low pJNK^T183/Y185^ activity in the TCGA breast cancer cohort [25, 30], where JNK phosphorylation was, on average, 37.5% lower in ER+ breast cancers with a *MAP2K7* homozygous or heterozygous loss compared to those cancers with no loss of *MAP2K7* (diploid, amplified or gain status) (p<0.0002; Figure 2F).

Since the JNK cascade is activated in response to a range of stress-inducing stimuli [49], we assessed whether MKK7 mediates stress-induced JNK signalling in ER+ breast cancer cells. Using anisomycin, an agent that inhibits protein translation to produce ribotoxic stress [50], JNK phosphorylation was rapidly induced within 30 minutes, and sustained for 90 minutes in MCF-7 pLenti cells (Figure 2G; Supplementary Figure 2I). In contrast, MCF-7 *MAP2K7*^-/-^ cells displayed markedly attenuated JNK phosphorylation within 30 minutes of anisomycin exposure and pJNK^T183/Y185^ levels returned to basal level by 60 minutes (Figure 2G). Anisomycin exposure in T-47D pLenti cells also led to a profound induction of JNK phosphorylation that was attenuated in T-47D *MAP2K7*^-/-^ cells between 30-60 minutes (Figure 2H; Supplementary Figure 2J). In summary, the loss of *MAP2K7* prevented a sustained stress-mediated induction of JNK phosphorylation in MCF-7 cells, while T-47D cells had a dampened response to persistent stress-activation.

One of the genes that encode JNK, *MAPK9*, was also a gene of interest from the CRISPR screen (Supplementary Table 1 and 2). *MAPK8* (encoding JNK1) and *MAPK9* (encoding JNK2) are similar genes that are functionally indistinguishable. To discern the effects of the individual genes, we generated *MAPK8* and *MAPK9* CRISPR/Cas9 knockouts in MCF-7 cells. After establishing that these cell lines exhibited almost complete knockout of their respective JNK protein (Figure 2I, J), we examined the effect of *MAPK8* and *MAPK9* loss on JNK phosphorylation. Loss of *MAPK8* and *MAPK9* led to a reduction in peak JNK phosphorylation levels 30 minutes following anisomycin exposure compared to pLenti cells, with a more profound decrease in *MAPK9*^-/-^ compared to *MAPK8*^-/-^ cell lines (Figure 2I, K; Supplementary Figure 2K).

Finally, we examined whether JNK phosphorylation was altered in association with *MAPK8* or *MAPK9* copy number loss using data from the TCGA cohort [25, 30]. ER+ breast cancers with *MAPK8* deletions had on average 21.6% lower JNK activity, whereas cancers with *MAPK9* deletion had 26.9% lower JNK phosphorylation than those without *MAPK9* deletions (Figure 2L). Overall, *MAPK8* and *MAPK9* deletions were associated with a loss of JNK phosphorylation but did not have as profound an effect as *MAP2K7* loss, either *in vitro* or in patient data.

### Treatment with combination endocrine therapy and CDK4/6 inhibition alters MKK7 and JNK phosphorylation

We next examined the response of *MAP2K7*^-/-^ cell lines to endocrine therapy and CDK4/6 inhibition. We exposed pLenti and *MAP2K7*^-/-^ MCF-7 and T-47D cells to endocrine therapy (tamoxifen or fulvestrant) plus palbociclib to determine if MKK7 and pJNK^T183/Y185^ were regulated by the drug combinations (Figure 3A, D). Tamoxifen plus palbociclib treatment partially reduced MKK7 expression in MCF-7 pLenti cells (Figure 3B), while treatment with either tamoxifen or fulvestrant plus palbociclib reduced MKK7 protein expression in T-47D pLenti cells (Figure 3E). MKK7 expression remained undetectable in *MAP2K7*^-/-^ cells, and *MAP2K7* mRNA expression was also unaltered with treatment (Figure 3B, E; Supplementary Figure 3A, B). pJNK^T183/Y185^ was significantly upregulated in MCF-7 pLenti cells exposed to both tamoxifen or fulvestrant plus palbociclib (Figure 3C, F). However, only tamoxifen plus palbociclib, and not fulvestrant plus palbociclib treatment led to an increase in JNK activity in T-47D pLenti cells. As before, the level of pJNK^T183/Y185^ was negligible to low in *MAP2K7* knockout MCF-7 and T-47D cell lines, and not altered with drug treatment (Figure 3C, F). In summary, ER+ breast cancer cells exhibit treatment-depending reductions in MKK7 and pJNK^T183/Y185^ expression.

**Figure 3:**
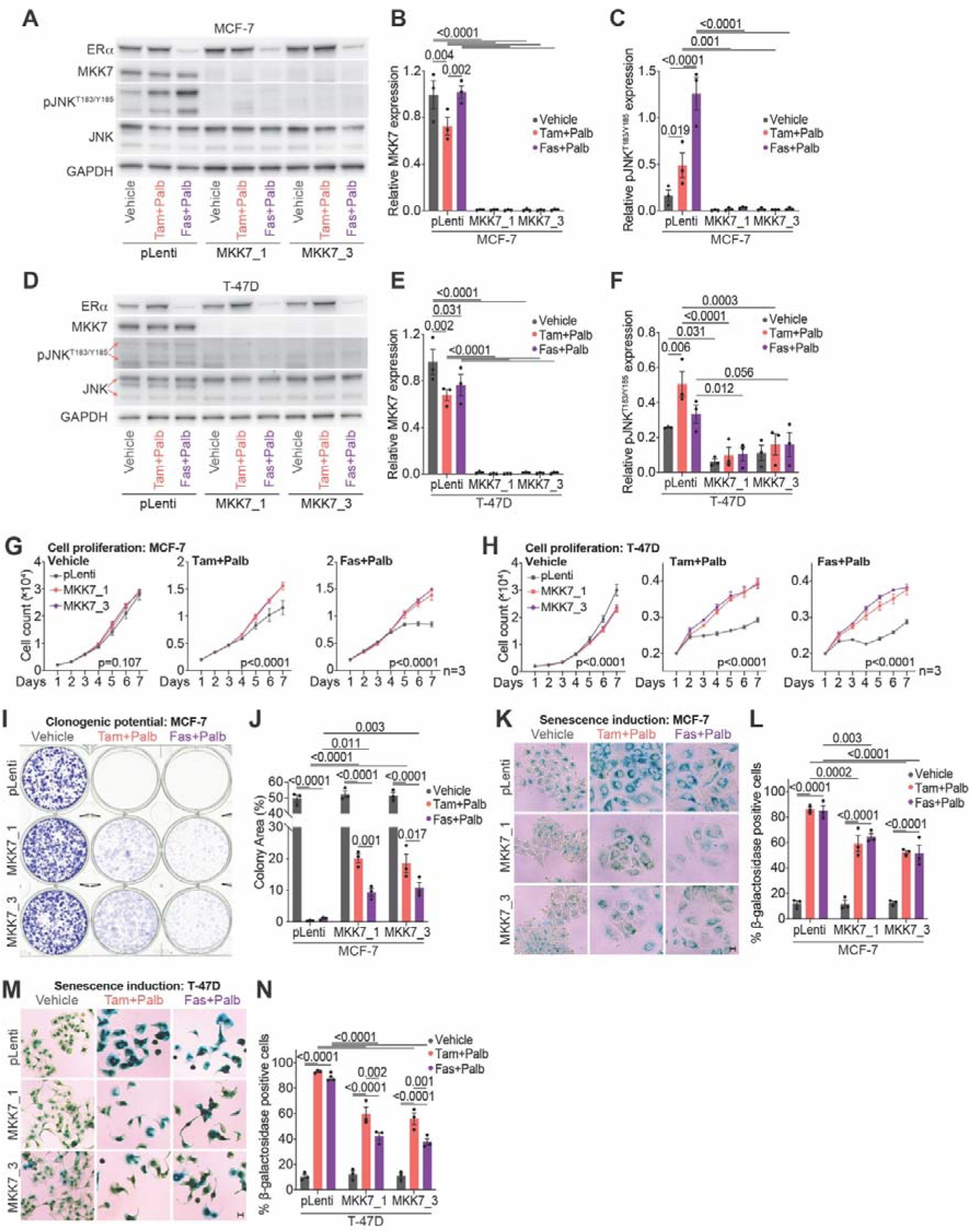
*MAP2K7* loss prevents growth arrest and senescence induction following endocrine therapy + CDK4/6 inhibitor treatment. **(A)** Representative Western blot of ERα, MKK7, pJNK^T183/Y185^, JNK and GAPDH in MCF-7 pLenti and *MAP2K7*^-/-^ (MKK7_1 and MKK7_3) cells treated with 500 nM tamoxifen + 250 nM palbociclib (tam+palb), 25 nM fulvestrant + 125 nM palbociclib (fas+palb) or vehicle (Absolute ethanol (EtOH) and tetrahydrofuran (THF)) for 48 hours (full length blots in Supplementary Figure 3B). **(B)** Quantitation of MKK7 expression by densitometry. Band intensity was normalised to GAPDH. Data analysed by two-way ANOVA with Tukey’s multiple comparisons test. **(C)** Quantitation of pJNK^T183/Y185^ activity by densitometry and normalised to GAPDH. Data analysed by two-way ANOVA with Tukey’s multiple comparisons test. **(D)** Representative Western blot of ERα, MKK7, pJNK^T183/Y185^, JNK and GAPDH in T-47D pLenti and *MAP2K7*^-/-^ (MKK7_1 and MKK7_3) cells treated with 500 nM tamoxifen + 250 nM palbociclib (tam+palb), 25 nM fulvestrant + 125 nM palbociclib (fas+palb) or vehicle (EtOH and THF) for 48 hours (full length blots in Supplementary Figure 3D). **(E)** Quantitation of MKK7 expression by densitometry and normalised to GAPDH. Data analysed by two-way ANOVA with Tukey’s multiple comparisons test. **(F)** Quantitation of pJNK^T183/Y185^ activity by densitometry and normalised to GAPDH. Data analysed by two-way ANOVA with Tukey’s multiple comparisons test. **(G)** MCF-7 pLenti and *MAP2K7*^-/-^ (MKK7_1 and MKK7_3) cells treated with 500 nM tamoxifen + 250 nM palbociclib (tam+palb), 25 nM fulvestrant + 125 nM palbociclib (fas+palb) or vehicle (EtOH and THF) and analysed by time-lapse microscopy using an IncuCyte ZOOM over 7 days. Cell count was determined by red fluorescent count, and data analysed by two-way ANOVA with Sidak’s multiple comparisons test. Experiment was performed in triplicate. **(H)** T-47D pLenti and *MAP2K7*^-/-^ (MKK7_1 and MKK7_3) cells treated with 500 nM tamoxifen + 250 nM palbociclib (tam+palb), 25 nM fulvestrant + 125 nM palbociclib (fas+palb) or vehicle (EtOH and THF) and analysed by time-lapse microscopy using an IncuCyte ZOOM over 7 days. Cell count was determined by red fluorescent count, and data analysed by two-way ANOVA with Sidak’s multiple comparisons test. Experiment was performed in triplicate. **(I)** Representative colony formations from MCF-7 pLenti and *MAP2K7*^-/-^ (MKK7_1 and MKK7_3) cells treated with 500 nM tamoxifen + 250 nM palbociclib (tam+palb), 25 nM fulvestrant + 125 nM palbociclib (fas+palb) or vehicle (EtOH and THF) for 3 weeks, with colony-formation detected with 0.1 – 0.5% crystal violet stain. **(J)** Colony formation was quantitated using ImageJ. Data analysed by two-way ANOVA with Tukey’s multiple comparisons test. **(K)** Representative brightfield images of MCF-7 pLenti and *MAP2K7*^-/-^ (MKK7_1 and MKK7_3) cells treated with 500 nM tamoxifen + 250 nM palbociclib (tam+palb), 25 nM fulvestrant + 125 nM palbociclib (fas+palb) or vehicle (EtOH and THF) for 72h and stained with senescence-associated β-galactosidase. **(L)** Quantification of MCF-7 pLenti and *MAP2K7*^-/-^ cells staining positive for senescence-associated β-galactosidase from (K). Data analysed by two-way ANOVA with Tukey’s multiple comparisons test. Scale bar = 100 µm. **(M)** Representative brightfield images of T-47D pLenti and *MAP2K7*^-/-^ (MKK7_1 and MKK7_3) cells treated with 500 nM tamoxifen + 250 nM palbociclib (tam+palb), 25 nM fulvestrant + 125 nM palbociclib (fas+palb) or vehicle (EtOH and THF) for 72h and stained with senescence-associated β-galactosidase. **(N)** Quantification of T-47D pLenti and *MAP2K7*^-/-^ cells staining positive for senescence-associated β-galactosidase from (M). Data analysed by two-way ANOVA with Tukey’s multiple comparisons test. Scale bar = 100 µm.

As we had identified that *MAP2K7* sgRNAs were particularly enriched in cells resistant to a therapeutic combination with anti-estrogens, we aimed to assess whether *MAP2K7* knockout had an effect on ERα expression and the ability of cells to respond to anti-estrogen therapies. MCF-7 and T-47D *MAP2K7*^-/-^ cells showed similar basal expression of ERα when compared to pLenti cells (Figure 3A, D; Supplementary Figure 3C-F). Fulvestrant treatment reduced ERα expression >3-fold compared to vehicle treatment in both cell lines, as expected with its mode of action (Supplementary Figure 3C, E).

### *MAP2K7* loss promotes proliferation and cell survival following treatment with combination endocrine therapy and CDK4/6 inhibition

Given that *MAP2K7* loss was associated with cell survival in our CRISPR/Cas9 screen, we explored the role of MKK7 in mediating both acute cell cycle regulation, and long-term proliferation and survival in cells treated with endocrine therapy plus palbociclib. We first examined whether *MAP2K7*^-/-^ cells failed to undergo complete cell cycle arrest when treated with endocrine therapy plus palbociclib for 48 hours. All cell lines underwent similar cell cycle arrest, regardless of *MAP2K7* knockout (Supplementary Figure 3G-I).

Next, we examined whether there were differences in cell number over an extended period using time-lapse microscopy for 7 days. MCF-7 *MAP2K7*^-/-^ cells showed similar basal rates of cell proliferation as vehicle-treated pLenti cells (Figure 3G; left panel), while T-47D pLenti cells proliferated slightly faster compared to *MAP2K7*^-/-^ cells (Figure 3H; left panel). In contrast, *MAP2K7*^-/-^ cells treated with the combination of endocrine therapy and palbociclib exhibited a significantly higher growth rate compared to pLenti cells (Figure 3G, H; middle and right panels).

We then assessed cell survival of *MAP2K7*^-/-^ cells using colony formation assays conducted over 3 weeks. MCF-7 *MAP2K7*^-/-^ cells treated with tamoxifen or fulvestrant plus palbociclib formed significantly more colonies compared to pLenti-treated cells (Figure 3I, J), indicating innate insensitivity to endocrine therapies and CDK4/6 inhibition. T-47D cells treated over the same timeframe did not produce sufficient colonies for analysis.

Given the reduced anisomycin-induced pJNK^T183/Y185^ activation in *MAPK8*^-/-^ and *MAPK9*^-/-^ MCF-7 cells (Figure 2I, K), we sought to evaluate whether these cells exhibited an altered response to endocrine therapy plus palbociclib treatment. To assess this, we measured changes to cell proliferation over 7 days using an IncuCyte assay. pLenti vehicle-treated cells proliferated significantly faster than *MAPK8*^-/-^ and *MAPK9*^-/-^ cells (p=0.0006; Supplementary Figure 3J; left panel). However, knockout of *MAPK8* or *MAPK9* did not alter growth rates of cells treated with endocrine therapy plus palbociclib compared to pLenti cells (Supplementary Figure 3J; middle and right panels). Overall, *MAP2K7*, but not *MAPK8* or *MAPK9* loss was associated with increased growth rates and cell survival following treatment with combination endocrine therapy and palbociclib treatment.

### *MAP2K7* knockout cells are refractory to senescence when treated with combination endocrine therapy and CDK4/6 inhibition

Since the induction of senescence is a major component of the cytostatic effect of endocrine therapy and CDK4/6 inhibitors [51, 52], we evaluated the impact of MKK7 on senescence response. We analysed MCF-7 and T-47D *MAP2K7*^-/-^ cells for senescence by detecting the accumulation of senescence-associated β-galactosidase. Combination therapy led to a 20- 50% decrease in the number of *MAP2K7*^-/-^ knockout cells staining positive for β- galactosidase compared to pLenti cells in both MCF-7 and T-47D cell lines (Figure 3K-N).

### *MAP2K7* loss diminishes the response to endocrine therapy and CDK4/6 inhibition via loss of activator protein-1 (AP-1) transcription factors

Given that activation of the JNK signalling cascade regulates apoptotic signalling [49], we initially hypothesised that the pro-apoptotic function of JNK signalling would be diminished following *MAP2K7* knockout in the tumour suppressive ER+ breast cancer context. Characterisation of BCL-2 pro- and anti-apoptotic proteins in pLenti and *MAP2K7*^-/-^ cells demonstrated no changes to the basal expression of proteins, except for a marginal increase in the expression of the pro-survival protein, BCL-2, in T-47D *MAP2K7*^-/-^ cells (Supplementary Figure 4A, B). Next, we used the BCL-2 inhibitor venetoclax, which has been reported to activate JNK expression [53, 54], to evaluate apoptotic induction through annexin V (Supplementary Figure 4C-E). There were no changes to venetoclax-induced apoptosis in *MAP2K7*^-/-^ compared to pLenti cells, and inconsistent increases in apoptosis induction between the MCF-7 *MAP2K7* sgRNAs (Supplementary Figure 4C, D). Consequently, we undertook an unbiased approach, performing RNAseq on *MAP2K7* knockout cells to determine how they responded differently to endocrine therapy plus palbociclib treatment.

pLenti and MKK7_3 MCF-7 and T47D cells were exposed to vehicle, tamoxifen plus palbociclib or fulvestrant plus palbociclib, and analysed by RNAseq (Figure 4A). MDS was applied to RNAseq data to visualise the relationships between samples based on gene expression profiles. MDS analysis showed that exposure to drug combinations led to significant changes in gene expression in dimension 1 that was dependent on drug treatment in both pLenti and *MAP2K7*^-/-^ cells, with treatment groups clustering closely together (Figure 4B). Interestingly, we observed that *MAP2K7*^-/-^ cells showed a diminished drug response in dimension 1 in each cell line and treatment pair: for example, MCF7 *MAP2K7*^-/-^ cells treated with fulvestrant plus palbociclib showed a 15% reduction in change in PC dimension 1 compared to pLenti cells exposed to fulvestrant plus palbociclib (Figure 4B). We then examined the effect of *MAP2K7* loss on fulvestrant plus palbociclib treatment by mapping the correlation between drug-induced gene regulation in MCF-7 pLenti and *MAP2K7*^-/-^ cells (Figure 4C). Gene regulation by fulvestrant plus palbociclib showed a significant linear correlation between pLenti and *MAP2K7*^-/-^ cells. Remarkably, *MAP2K7*^-/-^ cells showed an attenuated response in the upregulation and downregulation of the majority of genes affected by combination therapy. This led to a significant shift from perfect correlation (p<0.00001) [55]. The same analysis on fulvestrant plus palbociclib regulation was performed in T-47D pLenti and *MAP2K7*^-/-^ cells, and tamoxifen plus palbociclib gene regulation in MCF-7 and T- 47D cells. In all cell lines and treatment pairs, we observed a loss of perfect correlation in gene regulation, with less downregulation of downregulated targets, and less upregulation of upregulated targets (Figure 4C). Overall, this confirmed that gene expression in response to drug treatment was being attenuated following *MAP2K7* knockout, rather than new gene expression programs being induced by *MAP2K7* loss.

**Figure 4:**
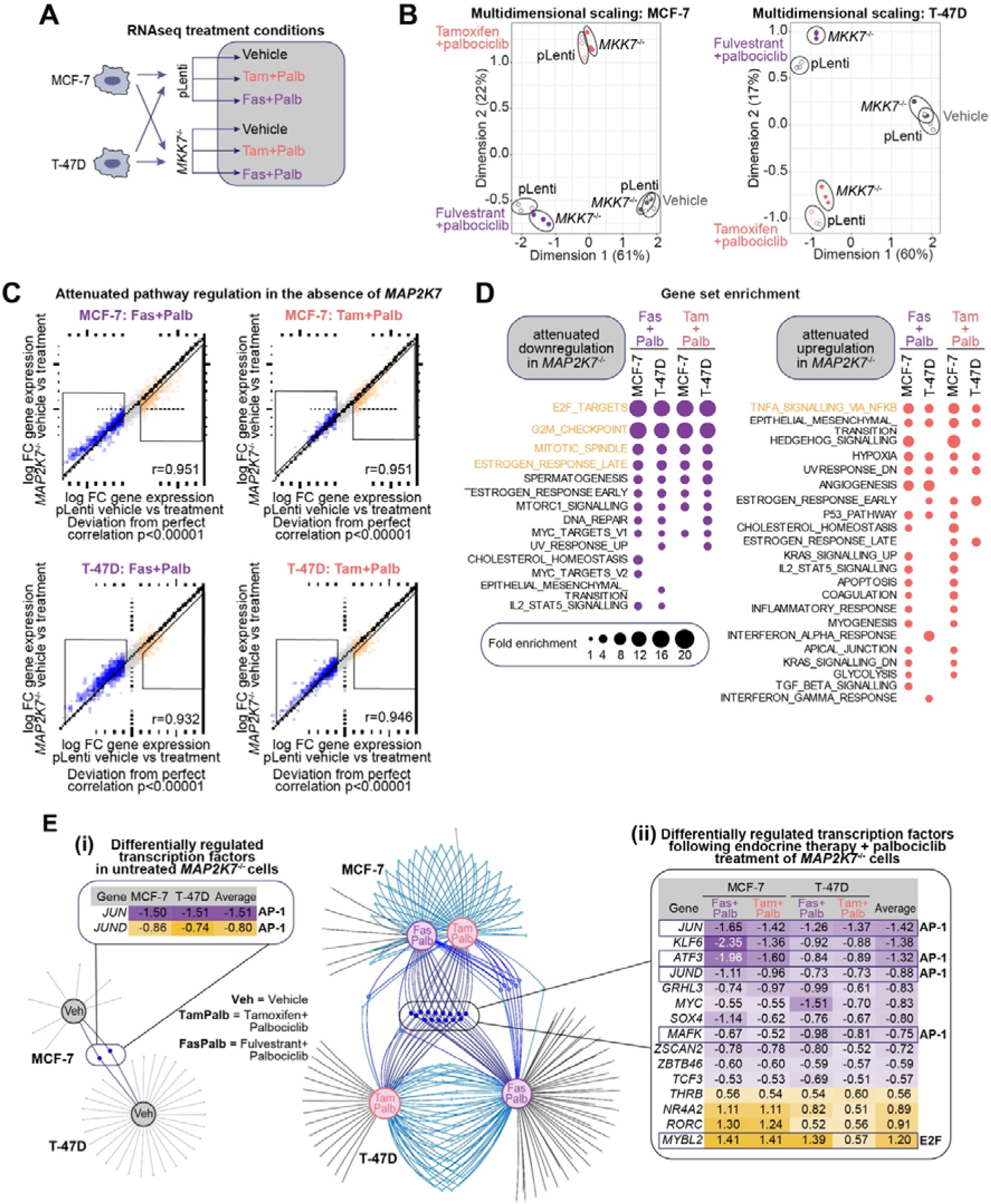
*MAP2K7* loss alters the transcriptional response to endocrine therapy + CDK4/6 inhibitor treatment. **(A)** Experimental schematic of pLenti and MKK7_3 cells (MCF-7 or T-47D) treated with 500 nM tamoxifen + 250 nM palbociclib (tam+palb), 25 nM fulvestrant + 125 nM palbociclib (fas+palb) or vehicle (Absolute ethanol (EtOH) and tetrahydrofuran (THF)) for 48 hours. RNA was collected and analysed by RNAseq. **(B)** Principal components analysis of RNAseq analysis of MCF-7 cells +/- treatments, and T-47D cells +/- treatments. **(C)** Relative change in gene expression between vehicle and treatment (tamoxifen + palbociclib or fulvestrant + palbociclib) of *MAP2K7*^-/-^ vs pLenti cells. Boxed regions are genes that are less downregulated, or less upregulated, in MKK7_3 cells compared to pLenti cells. Pearson’s correlation coefficient shown. **(D)** Analysis of gene set enrichment of hallmark gene sets from genes that are less downregulated (purple plots) or less upregulated (orange plots) with treatment in *MAP2K7*^-/-^ cells, in either MCF-7 or T-47D cell lines. Size of dot is fold enrichment of the signature, with enriched signatures with false discovery rate (FDR) of 0.05 shown. FDR values shown in Supplementary Table 3. **(E) (i)** Venn spider plots of transcription factors (TFs) commonly deregulated in *MAP2K7*^-/-^ MCF-7 and *MAP2K7*^-/-^ T- 47D cells that are untreated; Veh = vehicle; boxed region contains significantly altered TFs. **(ii)** Venn spider plots of TFs commonly deregulated in *MAP2K7*^-/-^ MCF-7 and T-47D cells that are treated with combination therapy; tam + palb = 500 nM tamoxifen + 250 nM palbociclib; or fas+palb = 25 nM fulvestrant + 125 nM palbociclib; boxed region contains significantly altered TFs.

We examined the attenuated pathways by performing GSEA of the genes that were less downregulated or upregulated in *MAP2K7*^-/-^ cells compared to pLenti cells. We used a cut-off of genes that are more than 1-fold downregulated in pLenti cells, which were then 0.25-fold less downregulated in *MAP2K7*^-/-^ cells, and examined enrichment of Hallmark gene sets. Using sets of downregulated genes, we identified that the most enriched datasets from each cell line and treatment condition were almost identical, including G2M_CHECKPOINT, E2F_TARGETS, MITOTIC_SPINDLE and ESTROGEN_RESPONSE_LATE (Figure 4D, left panel). These are pathways that are regulated downstream of anti-estrogen and CDK4/6 inhibitor treatment [5, 56]. Next, we examined upregulated gene signatures and found that the most consistently attenuated signature with *MAP2K7* loss was HALLMARK_TNFA_SIGNALLING_VIA_NFKB, which is a co-regulated pathway in estrogen signalling and in MAPK signalling [57] (Figure 4D).

Overall, *MAP2K7* loss attenuated transcriptional pathways controlling cell cycle response during drug treatment. Apart from TNFA_SIGNALLING_VIA_NFKB, we did not see changes to canonical outputs of MAPK signalling, which would be detected by signatures involving apoptosis, MAPK and stress. Consequently, we examined which transcription factors were most differentially regulated by loss of *MAP2K7*, both before and after drug treatment, to understand how transcriptional regulation is disrupted downstream of endocrine therapy and CDK4/6 inhibitor treatment. Firstly, we examined the difference in expression of transcription factors between vehicle-treated pLenti and *MAP2K7*^-/-^ cells, using a cut-off of log fold change >0.5. Only two transcription factors, *JUN* and *JUND*, were commonly downregulated with *MAP2K7* loss in MCF-7 and T-47D cell lines (Figure 4E(i)). *JUN* and *JUND* are both components of AP-1 transcriptional complexes that enable growth factor co-activation of estrogen signalling [58]. Furthermore, *JUN* was a hit in our CRISPR/Cas9 screen shown in Figure 1D and 1E.

Following treatment with endocrine therapy plus palbociclib, there was a greater number of transcription factors deregulated with *MAP2K7* loss in each condition (Figure 4E(ii)). There were 15 common deregulated transcription factors (downregulated transcription factors: *JUN*, *KLF6*, *ATF3*, *JUND*, *GRHL3*, *MYC*, *SOX4*, *MAFK*, *ZSCAN2*, *ZBTB46*, *TCF3*; upregulated transcription factors: *THRB*, *NR4A2*, *RORC*, *MYBL2*). Of these, four (*JUN*, *JUND*, *ATF3* and *MAFK*) were AP-1 component genes.

### cJUN transcription factor expression is depleted following *MAP2K7* loss

We next validated the relative protein and mRNA expression of AP-1 transcription factor genes in basal and treatment conditions of pLenti and *MAP2K7*^-/-^ MCF-7 and T-47D cells. Both MCF-7 and T-47D *MAP2K7*^-/-^ cells showed a marked reduction in p-cJUN^Ser63^ and total cJUN protein in basal conditions (Figure 5A-F). Treatment with tamoxifen or fulvestrant plus palbociclib led to a significant increase in the induction of p-cJUN^Ser63^ activity in both MCF-7 and T-47D cells, but in the absence of *MAP2K7,* the expression of *JUN* mRNA and p- cJUN^Ser63^/cJUN protein remained low or absent (Figure 5A-F; Supplementary Figure 5A, B; left panels). *JUND* mRNA and JUND protein were not altered at basal conditions in *MAP2K7*^-/-^ compared to pLenti cells, with the exception of a decrease in the expression of *JUND* in T-47D MKK7_3 cells (Figure 5A, D; Supplementary Figure 5A, B; middle panels; Supplementary Figure 5C, E). Only treatment with fulvestrant, and not tamoxifen plus palbociclib led to a decrease in JUND protein. However, this was not consistent between MCF-7 and T-47D cell lines and not unique to *MAP2K7*^-/-^ cells (Supplementary Figure 5C, E). Surprisingly, the inverse was seen with *JUND* mRNA expression, where treatment with endocrine therapy plus palbociclib increased expression (Supplementary Figure 5A, B; middle panels).

**Figure 5:**
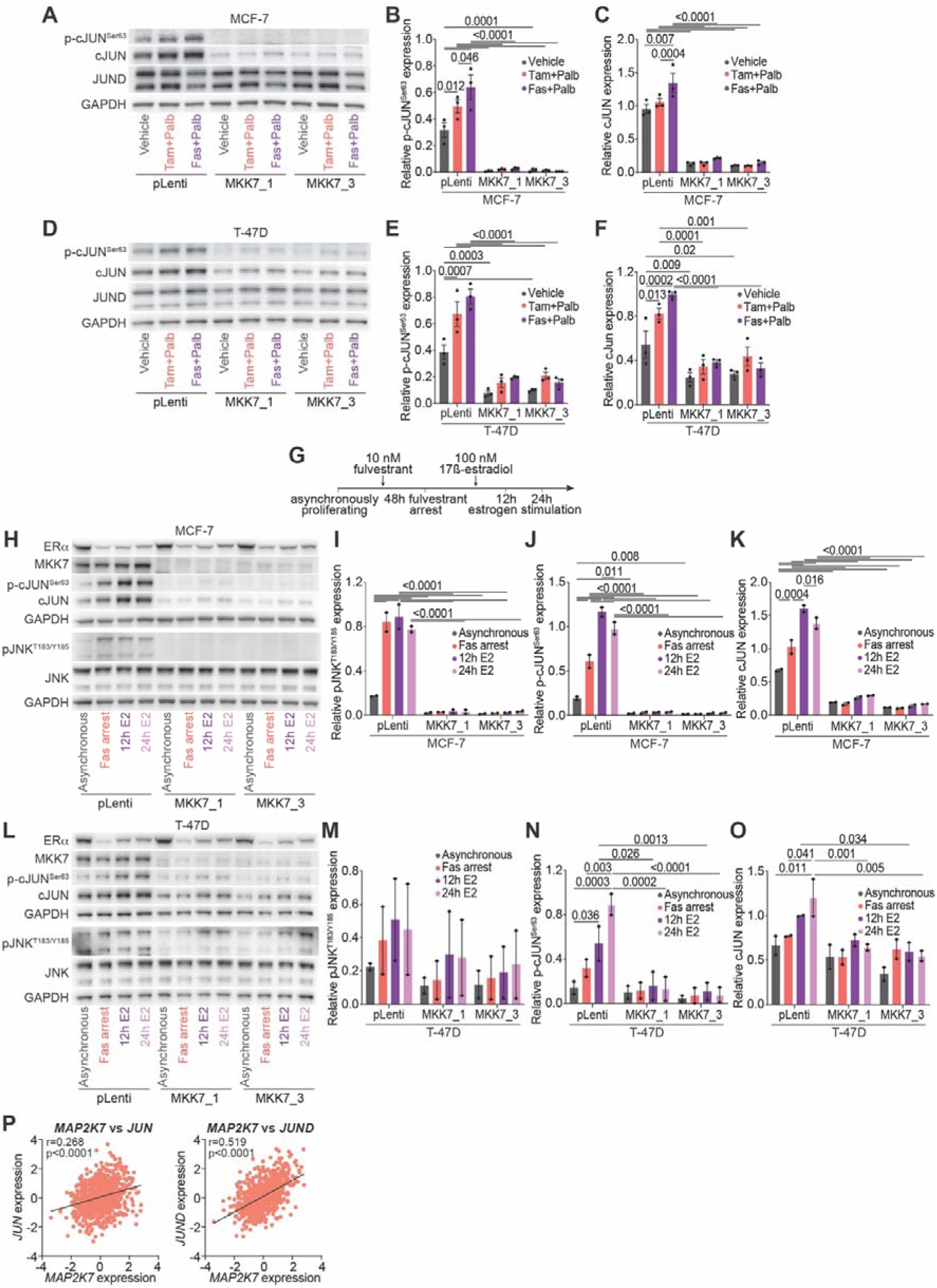
*MAP2K7* loss ameliorates AP-1 transcription factor signalling. **(A)** Representative Western blot of p-cJUN^Ser63^, cJUN, JUND and GAPDH in MCF-7 pLenti and *MAP2K7*^-/-^ (MKK7_1 and MKK7_3) cells treated with 500 nM tamoxifen + 250 nM palbociclib (tam+palb), 25 nM fulvestrant + 125 nM palbociclib (fas+palb) or vehicle (Absolute ethanol (EtOH) and tetrahydrofuran (THF)) for 48 hours (full length blots in Supplementary Figure 5D). **(B)** Quantitation of p-cJUN^Ser63^ activity by densitometry. Band intensity was normalised to GAPDH. Data analysed by two-way ANOVA with Tukey’s multiple comparisons test. **(C)** Quantitation of cJUN expression by densitometry and normalised to GAPDH. Data analysed by two-way ANOVA with Tukey’s multiple comparisons test. **(D)** Representative Western blot of p-cJUN^Ser63^, cJUN, JUND and GAPDH in T-47D pLenti and *MAP2K7*^-/-^ (MKK7_1 and MKK7_3) cells treated with 500 nM tamoxifen + 250 nM palbociclib (tam+palb), 25 nM fulvestrant + 125 nM palbociclib (fas+palb) or vehicle (EtOH and THF) for 48 hours (full length blots in Supplementary Figure 5F). **(E)** Quantitation of p-cJUN^Ser63^ activity by densitometry. Band intensity was normalised to GAPDH. Data analysed by two-way ANOVA with Tukey’s multiple comparisons test. **(F)** Quantitation of cJUN expression by densitometry and normalised to GAPDH. Data analysed by two-way ANOVA with Tukey’s multiple comparisons test. **(G)** Schematic of treatment to examine short-term fulvestrant-mediated arrest and release. MCF-7 and T-47D cells were treated with 10 nM fulvestrant for 48 hours to induce a cell cycle arrest. Cells were then stimulated to re-enter the cell cycle with 100 nM 17β-estradiol for 12 hours and 24 hours. **(H)** Representative Western blot of ERα, MKK7, p-cJUN^Ser63^, cJUN, pJNK^T183/Y185^, JNK and GAPDH in MCF-7 pLenti and *MAP2K7*^-/-^ (MKK7_1 and MKK7_3) cells treated with fulvestrant and estradiol (full length blots in Supplementary Figure 5I). **(I)** Quantitation of pJNK^T183/Y185^ activity by densitometry. Band intensity was normalised to GAPDH. Data analysed by two-way ANOVA with Tukey’s multiple comparisons test. **(J)** Quantitation of p-cJUN^Ser63^ activity by densitometry. Band intensity was normalised to GAPDH. Data analysed by two-way ANOVA with Tukey’s multiple comparisons test. **(K)** Quantitation of cJUN expression by densitometry. Band intensity was normalised to GAPDH. Data analysed by two-way ANOVA with Tukey’s multiple comparisons test. **(L)** Representative Western blot of ERα, MKK7, p-cJUN^Ser63^, cJUN, pJNK^T183/Y185^, JNK and GAPDH in T-47D pLenti and *MAP2K7*^-/-^ (MKK7_1 and MKK7_3) cells treated with fulvestrant and estradiol (full length blots in Supplementary Figure 5L). **(M)** Quantitation of pJNK^T183/Y185^ activity by densitometry. Band intensity was normalised to GAPDH. Data analysed by two-way ANOVA with Tukey’s multiple comparisons test. **(N)** Quantitation of p-cJUN^Ser63^ activity by densitometry. Band intensity was normalised to GAPDH. Data analysed by two-way ANOVA with Tukey’s multiple comparisons test. **(O)** Quantitation of cJUN expression by densitometry. Band intensity was normalised to GAPDH. Data analysed by two-way ANOVA with Tukey’s multiple comparisons test. **(P)** Expression of *JUN* and *JUND* in primary ER+ breast cancers from the TCGA cohort compared to *MAP2K7* expression. Pearson’s correlation coefficient shown.

Two other AP-1 transcription factors were altered in RNAseq analysis: ATF3 and MAFK. ATF3 protein was not detectable by Western blot in MCF-7 and T-47D cells, and mRNA expression was only altered in MCF-7 not T-47D cells (Supplementary Figure 5A, B; right panels). *MAFK* was expressed at very low transcript levels in both MCF-7 and T-47D cells (Supplementary Table 4).

Overall, the loss of *MAP2K7* led to a severe depletion of the AP-1 transcription factor cJUN, which is a fundamental co-factor of the ER. Consequently, we examined whether *MAP2K7* loss altered cell cycle progression in the context of endocrine therapy treatment and release (Figure 5G). All cell lines underwent similar G_1_ phase cell cycle arrest with fulvestrant treatment, and estradiol stimulation after 12 and 24h hours did not alter cell cycle distribution of *MAP2K7^-/-^*\ compared to pLenti cell lines (Supplementary Figure 5G, H).

Since acute changes in cell cycle response were not observed, we examined whether *MAP2K7* loss altered JNK phosphorylation and JUN regulation following ER blockade and stimulation (Figure 5H, L). In MCF-7 pLenti cells, the addition of fulvestrant led to an increase in pJNK^T183/Y185^ activity, and this high pJNK^T183/Y185^ level was maintained in cells stimulated with estradiol (Figure 5H, I). Loss of *MAP2K7* led to a loss of pJNK^T183/Y185^ activity with fulvestrant treatment, and estrogen stimulation in MCF-7 cells (Figure 5H, I). In T-47D pLenti and *MAP2K7*^-/-^ cells, the phosphorylation of JNK was maintained in fulvestrant-arrested cells, and following removal of fulvestrant and stimulation with estradiol (Figure 5L, M). Fulvestrant-arrested MCF-7 and T-47D pLenti cells had an increase in p-cJUN^Ser63^ activity and total protein expression compared to untreated, which was further enhanced with estrogen stimulation (Figure 5H, J-L, N, O). MCF-7 *MAP2K7*^-/-^ cells lacked p-cJUN^Ser63^ activity despite treatment with fulvestrant and estrogen (Figure 5H, J). In contrast, estrogen stimulation of T- 47D *MAP2K7*^-/-^ cells led to a decrease in p-cJUN^Ser63^ activity compared to treated T-47D pLenti cells (Figure 5L, N). Thus, in this model system JNK and cJUN are phosphorylated by fulvestrant arrest and estrogen recue in pLenti cells, but levels are dramatically reduced with *MAP2K7* knockout.

Additionally, we assessed the dynamic change in ERα and MKK7 expression after fulvestrant arrest and estradiol stimulation (Supplementary Figure 5I-N). MCF-7 and T-47D pLenti and *MAP2K7*^-/-^ cells had a similar fluctuation in ERα expression upon treatment with fulvestrant and estradiol (Supplementary Figure 5J, M). MKK7 expression was not altered by fulvestrant or estradiol treatment in pLenti or *MAP2K7*^-/-^ MCF-7 or T-47D cells (Supplementary Figure 5K, N).

Finally, we examined the relationship between JNK pathway genes and *JUN*/*JUND* expression in primary ER+ breast cancers from the TCGA (Figure 5P, Supplementary Figure 5O-Q) [30]. *MAP2K7* was positively correlated with *JUN* and *JUND* expression, whereas *MAP2K4*, *MAPK8* and *MAPK9* were negatively correlated with *JUN* and *JUND* expression. Therefore, *MAP2K7* is a probable primary driver of *JUN* stability via the JNK pathway in ER+ breast cancer.

### Loss of JNK pathway genes and pJNK^Y185^ activity is common in ER+ breast cancer

Our CRISPR screen identified JNK pathway loss, or potential tumour suppressive action, in the context of treatment for advanced/metastatic ER+ breast cancer. Since the JNK pathway is associated with both oncogenic and tumour suppressive functions depending on the tumour type [48], we systematically examined patient cohorts to elucidate the relationship of the JNK pathway and its individual genes in ER+ breast cancer.

Firstly, we examined the expression and phosphorylation of JNK1 (*MAPK8*) and JNK2 (*MAPK9*) in ER+ breast cancer using UALCAN data [33]. In normal tissue and primary luminal ER+ breast cancers, JNK1 and JNK2 proteins are expressed at similar levels (Figure 6A). However, we observed that pJNK2^Y185^ but not pJNK1^Y185^ can occur at much lower levels, with pJNK2^Y185^ expressed significantly lower in primary ER+ breast cancers than normal tissues (Figure 6A). These data indicate that JNK pathway downregulation is common in primary ER+ breast cancer.

**Figure 6:**
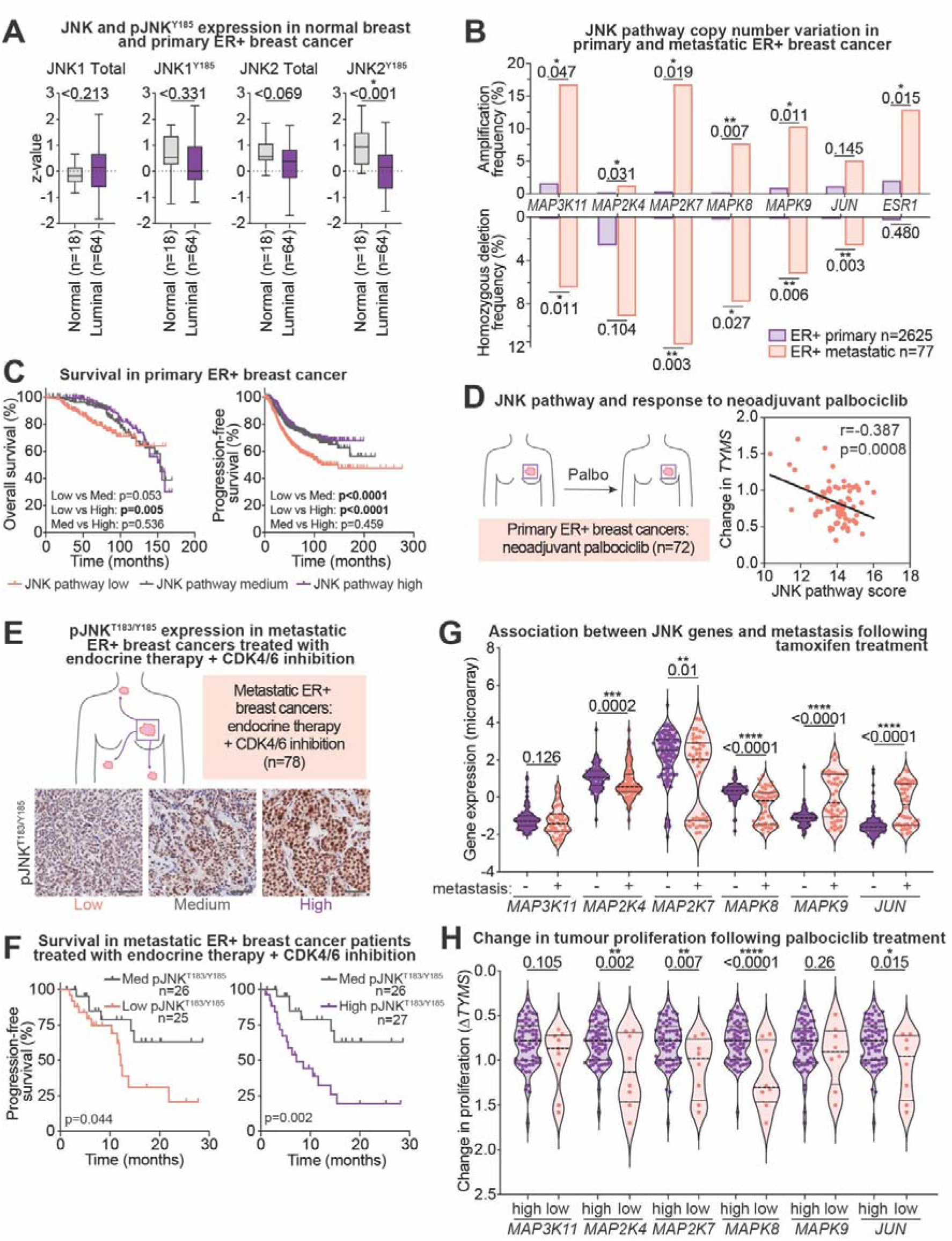
Depletion of JNK signalling in ER+ breast cancer, and in endocrine therapy/palbociclib resistance. **(A)** JNK1, JNK2, pJNK1^Y185^ (NP_001265477.1), pJNK2^Y185^ (NP_001128516.1) Clinical Proteomic Tumour Analysis Consortium data from the TCGA cohort, showing expression in normal breast tissue (n=18) and luminal breast cancers (n=64). Data analysed by unpaired two-tailed t-test. **(B)** Copy number status of *MAP3K11*, *MAP2K4*, *MAP2K7*, *MAPK8*, *MAPK9, JUN* and *ESR1* in primary (TCGA; n=808 and METABRIC; n=1817) and metastatic (Metastatic Breast Cancer Project; n=77) cohorts. Data analysed by unpaired two-tailed t-test. **(C)** Kaplan-Meier curves of the probability of overall survival and relapse-free survival in ER+ breast cancers comparing high, medium, and low tertiles of JNK pathway expression, where JNK pathway is *MAP2K7*, *MAPK8* and *MAPK9*. Kaplan-Meier analysis was performed on pooled breast cancer datasets using KMPlotter [73]. P-value calculated by log-rank (Mantel-Cox) test. **(D)** Correlation of JNK pathway with the anti-proliferative response of pre-operative palbociclib (POP) trial ER+ breast cancer patients [36]. Anti-proliferative response was determined by change in *TYMS* mRNA expression between initial biopsy and post-treatment (Δ*TYMS*) and correlated with JNK signalling (*MAP2K7*, *MAPK8* and *MAPK9*). Pearson’s correlation coefficient shown. **(E)** Immunohistochemistry analysis of advanced metastatic ER+ breast cancer cohort treated with endocrine therapy + CDK4/6 inhibition. Images of FFPE sections with high, medium, and low pJNK^T183/Y185^ activity. Scale bar = 100 µm. **(F)** Kaplan-Meier curves of survival in the endocrine + CDK4/6 inhibitor-treated cohort comparing pJNK^T183/Y185^ tertiles or low, medium and high pJNK^T183/Y185^. Significance determined by log-rank (Mantel-Cox) test. **(G)** mRNA expression of *MAP3K11*, *MAP2K4*, *MAP2K7*, *MAPK8*, *MAPK9* and *JUN* stratified into patients who did not develop metastatic disease (no metastasis; n=107) and patients who did (metastasis; n=48). Comparisons between metastatic and non-metastatic by Mann-Whitney unpaired two-tailed t-test. **(H)** Anti-proliferative response of POP trial ER+ breast cancer patients and association with individual JNK pathways genes. Patients with high (n=64) and low (n=8) JNK pathway gene expression (*MAP3K11*, *MAP2K4*, *MAP2K7*, *MAPK8*, *MAPK9* and *JUN*) were stratified based on *TYMS* anti-proliferative response. Comparisons between high and low by unpaired two-tailed t-test.

Next, we examined copy number alterations in JNK pathway genes, including those identified in our CRISPR/Cas9 screen, in primary and metastatic ER+ breast cancer genomic datasets [29–31]. JNK pathway genes were generally not altered in primary disease, except for low frequency deletion of *MAP2K4* (Figure 6B). There was a higher rate of loss of JNK pathway genes in metastatic disease with significantly more gene deletions in *MAP3K11, MAP2K7*, *MAPK8*, *MAPK9* and *JUN*, as well as significant amplification events, indicating the potential for the pathway to be both oncogenic and tumour suppressive in this context (Figure 6B). By contrast, *ESR1* amplification, which is a known oncogenic event in advanced ER+ breast cancer occurred exclusively in the metastatic setting, whereas *ESR1* deletion was notably absent (Figure 6B).

### Low expression of pJNK^T183/Y185^ correlates with poor survival in ER+ breast cancer patients treated with endocrine therapy and/or CDK4/6 inhibition

Next, we assessed the association between JNK pathway activation and prognosis in ER+ breast cancer. We used a composite gene expression score of *MAP2K7, MAPK8* and *MAPK9* since the other genes identified in our CRISPR/Cas9 screen have crosstalk with other signalling pathways. For example, *MAP3K11* also regulates ERK signalling [59] and *MAP2K4* regulates p38 [47]. Since JNK signalling can be both oncogenic and tumour suppressive, gene expression was split into tertiles. Low JNK signature was associated with worse overall survival compared to high JNK expression (p=0.005), and patients with low JNK expressing cancers had worse progression-free survival compared to patients with medium expression or high JNK pathway expression (low vs med: p<0.0001; low vs high: p<0.0001; Figure 6C).

Next, we examined the association of JNK signalling with clinical outcomes in patients treated with palbociclib. In the Pre-Operative Palbociclib (POP) trial, patients (n=72) underwent serial biopsies with intervening neoadjuvant palbociclib treatment [36]. Response to palbociclib was determined as relative changes in proliferation that occurred between the pre-treatment biopsy and post-treatment surgical sample, measured by proliferative marker, *TYMS*. JNK pathway expression (*MAP2K7*, *MAPK8*, *MAPK9*) was negatively correlated with proliferation, indicating that loss of JNK pathway expression was associated with poor response to palbociclib (Figure 6D).

We examined pJNK^T183/Y185^ activity by IHC in a second cohort of metastatic ER+ breast cancer patients treated with combined endocrine therapy and CDK4/6 inhibition (Supplementary Figure 6A) [23]. pJNK^T183/Y185^ activity exhibited predominantly nuclear staining which was quantified by H-score. pJNK^T183/Y185^ staining scores were stratified into low, medium and high pJNK^T183/Y185^ expressing tertiles to examine potential oncogenic and tumour suppressive roles for JNK in these samples (Figure 6E, Supplementary Figure 6B). Kaplan-Meier survival analysis showed significantly poorer progression-free survival in patients with low compared to medium pJNK^T183/Y185^ activity (p=0.044), and medium compared to high pJNK^T183/Y185^ activity (p=0.002) (Figure 6F). There was no significant difference between low and high pJNK^T183/Y185^-expressing cancers (p=0.214, graph not shown).

As we had identified that knockout of distinct JNK pathway genes were associated with poor therapeutic response in our CRISPR screen, we next examined the association of individual JNK pathway genes with metastasis-free survival following therapy. We investigated a cohort of breast cancers [34] where 48 of 155 patients (31%) developed distant metastases following 5 years of tamoxifen treatment (Figure 6G). Patients with low *MAP2K4*, *MAP2K7* or *MAPK8* expression or high *MAPK9* or *JUN* in their tumours had greater metastatic frequency following tamoxifen treatment (Figure 6G). We then examined individual target genes within the POP study, and their association with response. Low expression of *MAP2K4, MAP2K7, MAPK8* and *JUN* was significantly associated with poor response to palbociclib (Figure 6H).

## DISCUSSION

Our study employed an unbiased whole-genome approach to demonstrate that the JNK signalling cascade plays an important role in driving resistance to combination endocrine therapy and CDK4/6 inhibition in ER+ breast cancer. These findings provide new insights into the multi-faceted roles of JNK signalling and its implications in therapeutic resistance.

The role of JNK activation in ER+ breast cancer is particularly controversial. JNK activation has been shown to drive CDK4/6 inhibitor resistance [60] and is associated with poor prognosis in luminal B type breast cancers [61]. Models of endocrine therapy resistance are reported to have elevated JNK activity, suggesting that JNK signalling is oncogenic in this setting, and that JNK is activated in ER+ breast cancer cells with *ESR1* mutation [62]. Furthermore, JNK knockdown decreases ER+ breast cancer cell proliferation [63], an observation that we also see with *MAPK8* or *MAPK9* knockdown (Supplementary Figure 3J). Conversely, mouse models with inactivated JNK signalling have increased tumour incidence [64, 65]. This duality has led to a conflicting rationale to pursue both JNK inhibitors [62] and JNK activators [66] for ER+ breast cancer.

Due to this controversy, we examined both high and low expression of JNK pathway genes across patient cohorts. We observed that inactivation of JNK signalling, along with low expression of *MAP2K7* and *MAPK8,* were consistently associated with poor outcome or poor drug response in ER+ breast cancer following various treatment regimes. However, high pJNK^T183/Y185^ activity was also associated with poor outcome in endocrine therapy and CDK4/6 inhibitor-treated patients, and high expression of *MAPK9* and *JUN* associated with increased metastasis in a tamoxifen-treated cohort. These findings underscore the nuanced role of JNK signalling in ER+ breast cancer, where any deviation from normal/medium levels of JNK signalling, be it hyperactivation or suppression was associated with poor outcomes in ER+ breast cancer.

Amplification, deletion and mutation of JNK pathway components, particularly in the metastatic setting, suggest that polyclonal disease may involve simultaneous activation and inactivation of the pathway within individuals. MKK7, a primary kinase in this pathway, emerged as a critical regulator in our CRISPR screen. Interestingly, *MAP2K7* has been previously identified as a hit in screens for tamoxifen and fulvestrant resistance [67, 68], though never characterised. Our findings suggest that depletion of JNK signalling, particularly through *MAP2K7* loss, is a key element of resistance to endocrine therapy and CDK4/6 inhibition. *MAP2K7* knockout reduced sensitivity to endocrine therapies and palbociclib in both short- and long-term assays, suggesting that its role extends beyond canonical JNK pathway functions, such as apoptosis induction [69], to include modulation of stress signalling and senescence.

The interplay between JNK and ER signalling may further illuminate its role in endocrine therapy and CDK4/6 inhibitor resistance. Mechanistically, the depletion of JNK signalling appears intrinsically linked to anti-estrogen and estrogen function. JNK1 has been implicated in directly binding promoters in an AP-1/*ESR1* complex, moderating ER-directed transcription [63]. Although *MAPK8* and *MAPK9* knockout did not have profound effects in our models, functional substitution within the pathway and the dominant role of *MAP2K7* may explain this observation. JUN, a downstream effector of JNK, has also been implicated as an oncogene in ER+ breast cancer, reinforcing the complexity of JNK’s role in modulating ER signalling and therapy resistance [70].

Our current understanding of markers of combination resistance is largely based on patient data emerging from clinical trials [36, 39, 40, 43, 71, 72]. However, these studies are constrained by the limited scope of targets analysed, which usually entail gene panels of known or suspected cancer drivers. Notably, the JNK pathway is not represented on these panels apart from *MAP2K4*. Concerningly, the absence of JNK pathway components in genomic panels could negatively impact patient outcomes when considering therapeutic sequencing before and after CDK4/6 inhibitor use, particularly given the context-dependent effects of the JNK cascade.

## CONCLUSIONS

We have identified a subset of patients’ refractory to combination endocrine therapy and CDK4/6 inhibition exhibit defective JNK pathway signalling, and loss of this pathway alters drug-induced stress signalling and senescence responses. We provide the rationale to screen advanced ER+ breast cancers for pJNK^T183/Y185^ and *MAP2K7* expression and caution against the use of JNK inhibitors in this setting.

## Abbreviations

AP-1: activator protein-1
CDK4/6: cyclin-dependent kinase 4/6
DEGs: differentially expressed genes
ER: estrogen receptor
EtOH: Absolute ethanol
FBS: fetal bovine serum
FDR: false discovery rate
FFPE: formalin-fixed paraffin-embedded
gDNA: genomic DNA
gRNAs: guide RNAs
GSEA: gene set enrichment analysis
IC_50_: half-maximal inhibitory concentration
IHC: immunohistochemistry
JNK: c-Jun N-terminal kinase
MBCP: Metastatic Breast Cancer Project
MDS: multidimensional scaling
METABRIC: Molecular Taxonomy of Breast Cancer International Consortium
MOI: multiplicity of infection
PC: Principal components
PCR: polymerase chain reaction
PI: propidium iodide
POP: Pre-Operative Palbociclib (POP) trial
PVDF: Polyvinylidene fluoride
SEM: standard error of the mean
sgRNAs: single guide RNA
TCGA: The Cancer Genome Atlas
VCFG: Victorian Centre for Functional Genomics

## Declarations

### Author contribution

Conceptualization: C.E.C; Investigation: S.A, C.S.L, K.J.F, C.E.W, Z.P, A.L.C, C.L.A; Methodology: S.A, C.L, K.J.F, D.R, I.N, K.J.S, C.E.C; Software: S.A, J.R; Formal analysis: S.A, C.L.A, E.K.A.M, H.J.D, C.E.C; Bioinformatics analysis: L.E; Resources: K.J.S, H.J.D; Writing – Original Draft: S.A, C.E.C; Writing – Review & Editing: S.A, T.E.H, C.E.C; Supervision: C.E.C; Funding acquisition: C.E.C.

## Acknowledgments

This study was supported by the following facilities at the Garvan Institute of Medical Research: Australian BioResource, Biological Testing Facility, Garvan Molecular Genetics, Garvan Sequencing Platform, Garvan-Weizmann Centre for Cellular Genomics Flow Facility, Tissue Culture Facility under Gillian Lehrbach and Ruth Lyons, and the Garvan Histology Core Facility under Anaiis Zaratzian. Next-generation sequencing for CRISPR/Cas9 screens were performed at the Molecular Genomics Core facility at Peter MacCallum Cancer Centre. We would like to thank Dr Twishi Gulati and the Victorian Centre for Function Genomics for their expertise and support with CRISPR/Cas9 screening.

## SUPPLEMENTARY FIGURES

**Supplementary Figure 1: Quality control from the CRISPR/Cas9 screen and MCF-7 dose response curves**

**(A)** Total reads, both mapped and unmapped, across 2 biological CRISPR/Cas9 screens. **(B)** Gini index (a measure of inequality, with 1 being the most unequal) compared to the starting cell population (T_0_) and between replicates. Data in **(A** and **B)** were analysed using MAGeCK-VISPR [12]. **(C)** Dose response curve of MCF-7 cells treated for 5 days with tamoxifen. IC_50_ values were determined by log transforming, normalising, and plotting data with a nonlinear fit of least square analysis. Experiment was performed in quadruplicate**. (D)** Dose response curve of MCF-7 cells treated for 5 days with palbociclib. IC_50_ values were determined by log transforming, normalising, and plotting data with a nonlinear fit of least square analysis. Experiment was performed in triplicate. **(E)** Growth rates of MCF-7 cells treated for up to 90 days with 500 nM palbociclib (palb), 500 nM tamoxifen + 250 nM palbociclib (tam+palb), or vehicle (tetrahydrofuran). Dotted line indicates starting cell number.

**Supplementary Figure 2: Effect of *MAP2K7* knockout on MKK4 and JNK expression**

**(A)** Quantitation of MKK4 protein expression in MCF-7 pLenti and *MAP2K7*^-/-^ (MKK7_1 and MKK7_3) cells by densitometry. Band intensity was normalised to respective GAPDH. Data analysed by one-way ANOVA with Tukey’s multiple comparisons test. **(B)** Uncropped western blots for Figure 2A. Polyvinylidene fluoride (PVDF) membranes were sliced into smaller sections to incubate with primary antibodies to allow accurate comparison of protein expression within a single sample. Full sections of the sliced membranes from each cropped Western blot are shown. **(C)** Quantitation of MKK4 protein expression in T-47D pLenti and *MAP2K7*^-/-^ (MKK7_1 and MKK7_3) cells by densitometry. Band intensity was normalised to respective GAPDH. Data analysed by one-way ANOVA with Tukey’s multiple comparisons test. **(D)** Uncropped western blots for Figure 2B. PVDF membranes were sliced into smaller sections to incubate with primary antibodies to allow accurate comparison of protein expression within a single sample. Full sections of the sliced membranes from each cropped western blot are shown. **(E)** Quantitation of JNK protein expression in MCF-7 pLenti and *MAP2K7*^-/-^ (MKK7_1 and MKK7_3) cells by densitometry. Band intensity was normalised to respective GAPDH. Data analysed by one-way ANOVA with Tukey’s multiple comparisons test. **(F)** Uncropped western blots for Figure 2D. PVDF membranes were sliced into smaller sections to incubate with primary antibodies to allow accurate comparison of protein expression within a single sample. Full sections of the sliced membranes from each cropped Western blot are shown. **(G)** Quantitation of JNK protein expression in T-47D pLenti and *MAP2K7*^-/-^ (MKK7_1 and MKK7_3) cells by densitometry. Band intensity was normalised to respective GAPDH. Data analysed by one-way ANOVA with Tukey’s multiple comparisons test. **(H)** Uncropped Western blots for Figure 2E. PVDF membranes were sliced into smaller sections to incubate with primary antibodies to allow accurate comparison of protein expression within a single sample. Full sections of the sliced membranes from each cropped western blot are shown. **(I)** Uncropped Western blots for Figure 2G. PVDF membranes were sliced into smaller sections to incubate with primary antibodies to allow accurate comparison of protein expression within a single sample. Full sections of the sliced membranes from each cropped western blot are shown. **(J)** Uncropped Western blots for Figure 2H. PVDF membranes were sliced into smaller sections to incubate with primary antibodies to allow accurate comparison of protein expression within a single sample. Full sections of the sliced membranes from each cropped western blot are shown. **(K)** Uncropped Western blots for Figure 2I. PVDF membranes were sliced into smaller sections to incubate with primary antibodies to allow accurate comparison of protein expression within a single sample. Full sections of the sliced membranes from each cropped Western blot are shown.

**Supplementary Figure 3: Effect of *MAP2K7* knockout on cell cycle regulation, and *MAPK8 and MAPK9* knockout on proliferation**

**(A)** Quantitation of ERα protein expression in MCF-7 pLenti and *MAP2K7*^-/-^ (MKK7_1 and MKK7_3) cells treated with 500 nM tamoxifen + 250 nM palbociclib (tam+palb), 25 nM fulvestrant + 125 nM palbociclib (fas+palb) or vehicle (Absolute ethanol (EtOH) and tetrahydrofuran (THF)) for 48 hours by densitometry. Band intensity was normalised to respective GAPDH. Data analysed by Two-way ANOVA with Tukey’s multiple comparisons test. **(B)** Uncropped Western blots for Figure 3A. Polyvinylidene fluoride (PVDF) membranes were sliced into smaller sections to incubate with primary antibodies to allow accurate comparison of protein expression within a single sample. Full sections of the sliced membranes from each cropped Western blot are shown. **(C)** Quantitation of ERα protein expression in T-47D pLenti and *MAP2K7*^-/-^ (MKK7_1 and MKK7_3) cells treated with 500 nM tamoxifen + 250 nM palbociclib (tam+palb), 25 nM fulvestrant + 125 nM palbociclib (fas+palb) or vehicle (EtOH and THF) for 48 hours by densitometry. Band intensity was normalised to respective GAPDH. Data analysed by Two-way ANOVA with Tukey’s multiple comparisons test. **(D)** Uncropped Western blots for Figure 3D. PVDF membranes were sliced into smaller sections to incubate with primary antibodies to allow accurate comparison of protein expression within a single sample. Full sections of the sliced membranes from each cropped Western blot are shown. **(E)** Quantitation of *MAP2K7* mRNA in MCF-7 pLenti and *MAP2K7*^-/-^ (MKK7_1 and MKK7_3) cells treated with 500 nM tamoxifen + 250 nM palbociclib (tam+palb), 25 nM fulvestrant + 125 nM palbociclib (fas+palb) or vehicle (EtOH and THF) for 48 hours. Expression was normalised to *RPLP0* and data analysed by Two-way ANOVA with Tukey’s multiple comparisons test. **(F)** Quantitation of *MAP2K7* mRNA in T-47D pLenti and *MAP2K7*^-/-^ (MKK7_1 and MKK7_3) cells treated with 500 nM tamoxifen + 250 nM palbociclib (tam+palb), 25 nM fulvestrant + 125 nM palbociclib (fas+palb) or vehicle (EtOH and THF) for 48 hours. Expression was normalised to *RPLP0* and data analysed by Two-way ANOVA with Tukey’s multiple comparisons test. **(G)** Cell cycle distribution determined by propidium iodide staining and flow cytometry of MCF-7 pLenti and *MAP2K7*^-/-^ (MKK7_1 and MKK7_3) cells treated with 500 nM tamoxifen + 250 nM palbociclib (tam+palb), 25 nM fulvestrant + 125 nM palbociclib (fas+palb) or vehicle (EtOH and THF) for 48 hours. No significant changes in cell cycle distribution between pLenti and *MAP2K7*^-/-^ cells were observed when analysed with χ^2^ test. Experiment was performed in triplicate. **(H)** Cell cycle distribution determined by propidium iodide staining and flow cytometry of T-47D pLenti and *MAP2K7*^-/-^ (MKK7_1 and MKK7_3) cells treated with 500 nM tamoxifen + 250 nM palbociclib (tam+palb), 25 nM fulvestrant + 125 nM palbociclib (fas+palb) or vehicle (EtOH and THF) for 48 hours. No significant changes in cell cycle distribution between pLenti and *MAP2K7*^-/-^ cells were observed when analysed with χ^2^ test. Experiment was performed in triplicate. **(I)** Gating strategy for cell cycle analysis by flow cytometry as shown in (G) and (H). **(J)** MCF-7 pLenti, *MAPK8*^-/-^ and *MAPK9*^-/-^ cells treated with 500 nM tamoxifen + 250 nM palbociclib (tam+palb), 25 nM fulvestrant + 125 nM palbociclib (fas+palb) or vehicle (EtOH and THF) were analysed by time-lapse microscopy using an IncuCyte ZOOM over 7 days. Cell count was determined by red fluorescent count, and data analysed by two-way ANOVA with Sidak’s multiple comparisons test. Experiment was performed in triplicate.

**Supplementary Figure 4: *MAP2K7^-/-^* ER+ breast cancer cell lines do not show changes in basal levels of apoptotic proteins, or altered response to BCL-2 inhibition**

**(A)** Representative Western blot of MCL-1, PUMA, BIM, BCL-XL, BCL-2 and GAPDH in MCF-7 pLenti and *MAP2K7*^-/-^ (MKK7_1 and MKK7_3) cells. Quantitation of proteins by densitometry. Band intensity was normalised to GAPDH. Data analysed by one-way ANOVA with Tukey’s multiple comparisons test. **(B)** Representative western blot of MCL-1, PUMA, BIM, BCL-XL, BCL-2 and GAPDH in T-47D pLenti and *MAP2K7*^-/-^ (MKK7_1 and MKK7_3) cells. Quantitation of proteins by densitometry. Band intensity was normalised to GAPDH. Data analysed by one-way ANOVA with Tukey’s multiple comparisons test. **(C)** MCF-7 pLenti and *MAP2K7*^-/-^ (MKK7_1 and MKK7_3) cells treated with vehicle or 10 µM venetoclax for 24 hours were stained with FITC-conjugated Annexin V and propidium iodide (PI) and analysed by flow cytometry. Data was analysed using FlowJo. **(D)** T-47D pLenti and *MAP2K7*^-/-^ (MKK7_1 and MKK7_3) cells treated with vehicle or 10 µM venetoclax for 24 hours were stained with FITC-conjugated Annexin V and PI and analysed by flow cytometry. Data was analysed using FlowJo on duplicate data. **(E)** Gating strategy for Annexin V analysis by flow cytometry as shown in (C) and (D).

**Supplementary Figure 5: Estrogen induction and JUN/JUND expression with *MAP2K7* loss**

**(A)** Quantitation of *JUN, JUND* and *ATF3* mRNA in MCF-7 pLenti and *MAP2K7*^-/-^ (MKK7_1 and MKK7_3) cells treated with 500 nM tamoxifen + 250 nM palbociclib (tam+palb), 25 nM fulvestrant + 125 nM palbociclib (fas+palb) or vehicle (Absolute ethanol (EtOH) and tetrahydrofuran (THF)) for 48 hours. Expression was normalised to *RPLP0* and data analysed by Two-way ANOVA with Tukey’s multiple comparisons test. pLenti samples were run in duplicate for *JUN* mRNA. **(B)** Quantitation of *JUN, JUND* and *ATF3* mRNA in T-47D pLenti and *MAP2K7*^-/-^ (MKK7_1 and MKK7_3) cells treated with 500 nM tamoxifen + 250 nM palbociclib (tam+palb), 25 nM fulvestrant + 125 nM palbociclib (fas+palb) or vehicle (EtOH and THF) for 48 hours. Expression was normalised to *RPLP0* and data analysed by Two-way ANOVA with Tukey’s multiple comparisons test. pLenti samples were run in duplicate for *JUN* mRNA. **(C)** Quantitation of JUND protein expression in MCF-7 pLenti and *MAP2K7*^-/-^ (MKK7_1 and MKK7_3) cells treated with 500 nM tamoxifen + 250 nM palbociclib (tam+palb), 25 nM fulvestrant + 125 nM palbociclib (fas+palb) or vehicle (EtOH and THF) for 48 hours by densitometry. Band intensity was normalised to respective GAPDH. Data analysed by Two-way ANOVA with Tukey’s multiple comparisons test. **(D)** Uncropped Western blots for Figure 5A. Polyvinylidene fluoride (PVDF) membranes were sliced into smaller sections to incubate with primary antibodies to allow accurate comparison of protein expression within a single sample. Full sections of the sliced membranes from each cropped Western blot are shown. **(E)** Quantitation of JUND protein expression in T-47D pLenti and *MAP2K7*^-/-^ (MKK7_1 and MKK7_3) cells treated with 500 nM tamoxifen + 250 nM palbociclib (tam+palb), 25 nM fulvestrant + 125 nM palbociclib (fas+palb) or vehicle (EtOH and THF) for 48 hours by densitometry. Band intensity was normalised to respective GAPDH. Data analysed by Two-way ANOVA with Tukey’s multiple comparisons test. **(F)** Uncropped Western blots for Figure 5D. PVDF membranes were sliced into smaller sections to incubate with primary antibodies to allow accurate comparison of protein expression within a single sample. Full sections of the sliced membranes from each cropped western blot are shown. **(G)** Cell cycle distribution determined by propidium iodide staining and flow cytometry of MCF-7 pLenti and *MAP2K7*^-/-^ (MKK7_1 and MKK7_3) cells treated with 10 nM fulvestrant for 48 hours and stimulated with 100 nM 17β-estradiol for 12 hours and 24 hours. No significant changes in cell cycle distribution between pLenti and *MAP2K7*^-/-^ cells were observed when analysed with χ^2^ test. Experiment was performed in duplicate. **(H)** Cell cycle distribution determined by propidium iodide staining and flow cytometry of T-47D pLenti and *MAP2K7*^-/-^ (MKK7_1 and MKK7_3) cells treated with 10 nM fulvestrant for 48 hours and stimulated with 100 nM 17β-estradiol for 12 hours and 24 hours. No significant changes in cell cycle distribution between pLenti and *MAP2K7*^-/-^ cells were observed when analysed with χ^2^ test. Experiment was performed in singlet. **(I)** Uncropped Western blots for Figure 5H. PVDF membranes were sliced into smaller sections to incubate with primary antibodies to allow accurate comparison of protein expression within a single sample. Full sections of the sliced membranes from each cropped western blot are shown. **(J)** Quantitation of ERα protein expression in MCF-7 pLenti and *MAP2K7*^-/-^ (MKK7_1 and MKK7_3) cells treated with fulvestrant and estradiol. Band intensity was normalised to respective GAPDH. Data analysed by Two-way ANOVA with Tukey’s multiple comparisons test. **(K)** Quantitation of MKK7 protein expression in MCF-7 pLenti and *MAP2K7*^-/-^ (MKK7_1 and MKK7_3) cells treated with fulvestrant and estradiol. Band intensity was normalised to respective GAPDH. Data analysed by Two-way ANOVA with Tukey’s multiple comparisons test. **(L)** Uncropped Western blots for Figure 5L. PVDF membranes were sliced into smaller sections to incubate with primary antibodies to allow accurate comparison of protein expression within a single sample. Full sections of the sliced membranes from each cropped western blot are shown. **(M)** Quantitation of ERα protein expression in T-47D pLenti and *MAP2K7*^-/-^ (MKK7_1 and MKK7_3) cells treated with fulvestrant and estradiol. Band intensity was normalised to respective GAPDH. Data analysed by Two-way ANOVA with Tukey’s multiple comparisons test. **(N)** Quantitation of MKK7 protein expression in T-47D pLenti and *MAP2K7*^-/-^ (MKK7_1 and MKK7_3) cells treated with fulvestrant and estradiol. Band intensity was normalised to respective GAPDH. Data analysed by Two-way ANOVA with Tukey’s multiple comparisons test. **(O)** Expression of *JUN* and *JUND* in primary ER+ breast cancers from the TCGA cohort compared to *MAP2K4* expression. Pearson’s correlation coefficient shown. **(P)** Expression of *JUN* and *JUND* in primary ER+ breast cancers from the TCGA cohort compared to *MAPK8* expression. Pearson’s correlation coefficient shown. **(Q)** Expression of *JUN* and *JUND* in primary ER+ breast cancers from the TCGA cohort compared to *MAPK9* expression. Pearson’s correlation coefficient shown.

**Supplementary Figure 6: Patient cohort evaluation**

**(A)** Flow diagram showing the number of formalin-fixed paraffin-embedded samples from endocrine therapy + CDK4/6 inhibitor-treated ER+ patients that were stained with pJNK^T183/Y185^ antibody for immunohistochemistry (IHC) analysis. **(B)** The distribution of IHC tumour H-scores for pJNK^T183/Y185^ activity split into high, medium, and low.

## SUPPLEMENTARY TABLES

**Supplementary Table 1:** sgRNA hits from CRISPR/Cas9 screens

**Supplementary Table 2:** Common sgRNA hits from CRISPR/Cas9 screens

**Supplementary Table 3:** Transcription factors from RNAseq

**Supplementary Table 4:** RNAseq counts per million

