## Supplementary figures and images for "JNK pathway suppression drives resistance to combination endocrine therapy and CDK4/6 inhibition in ER+ breast cancer"

### Supplementary Figures

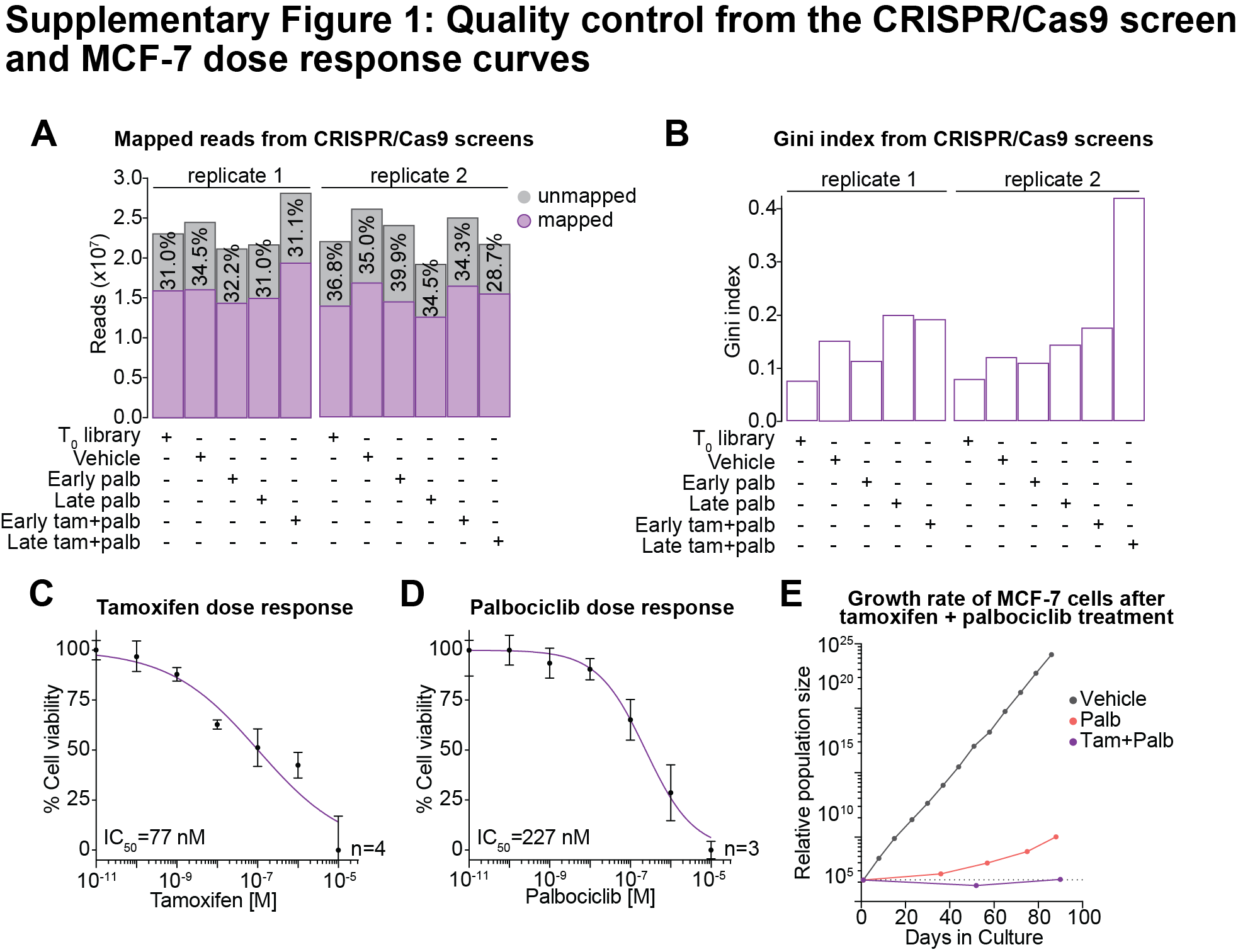

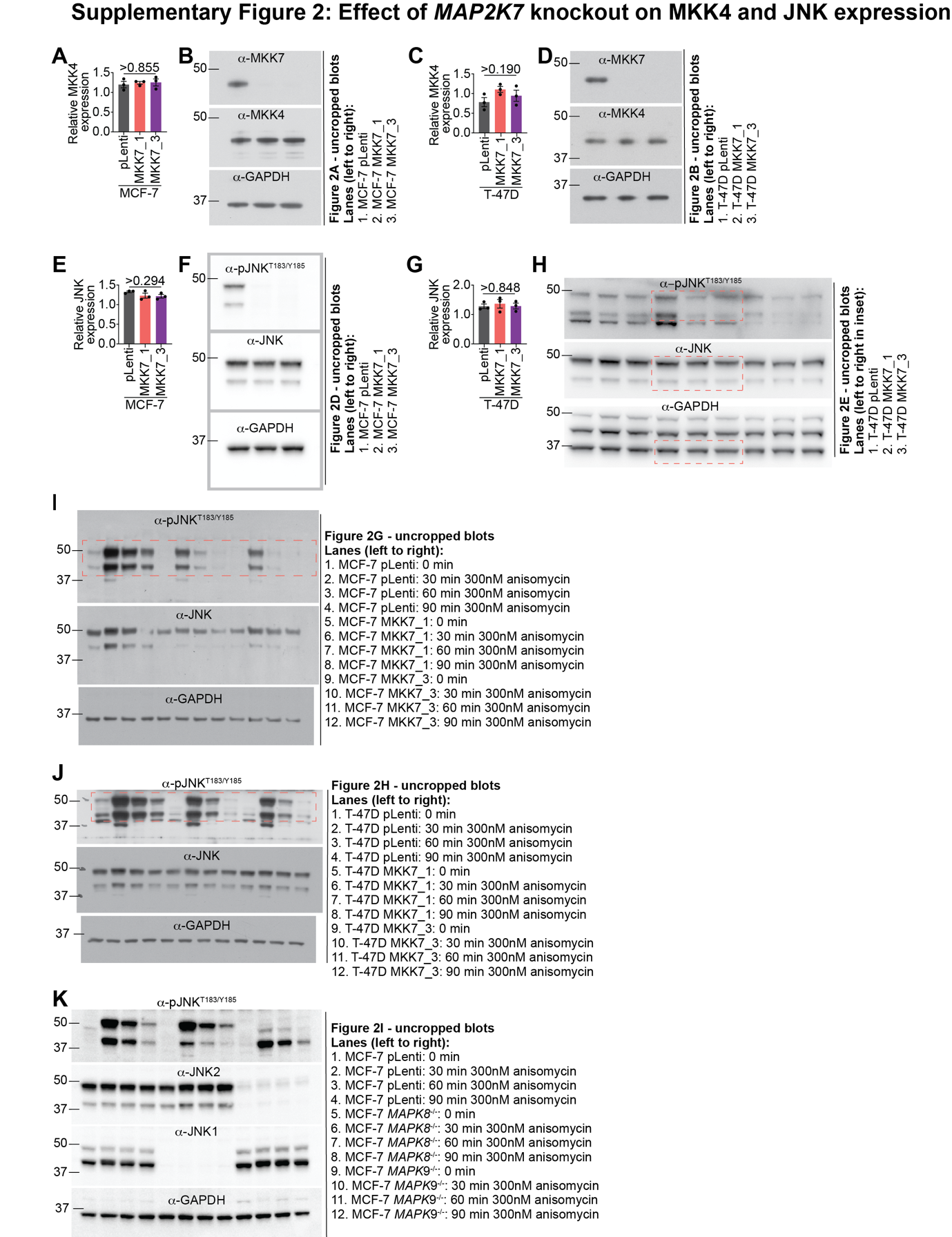

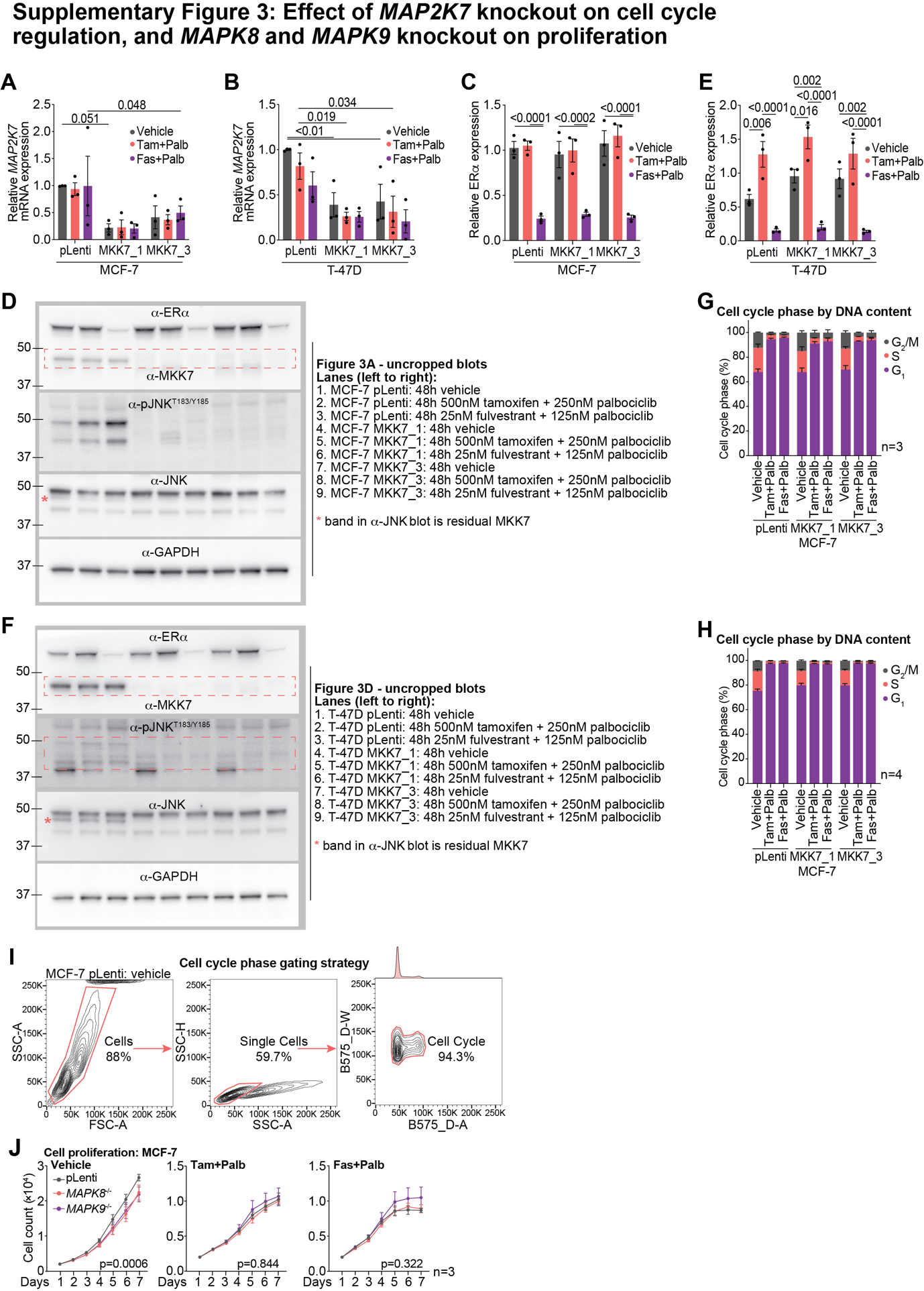

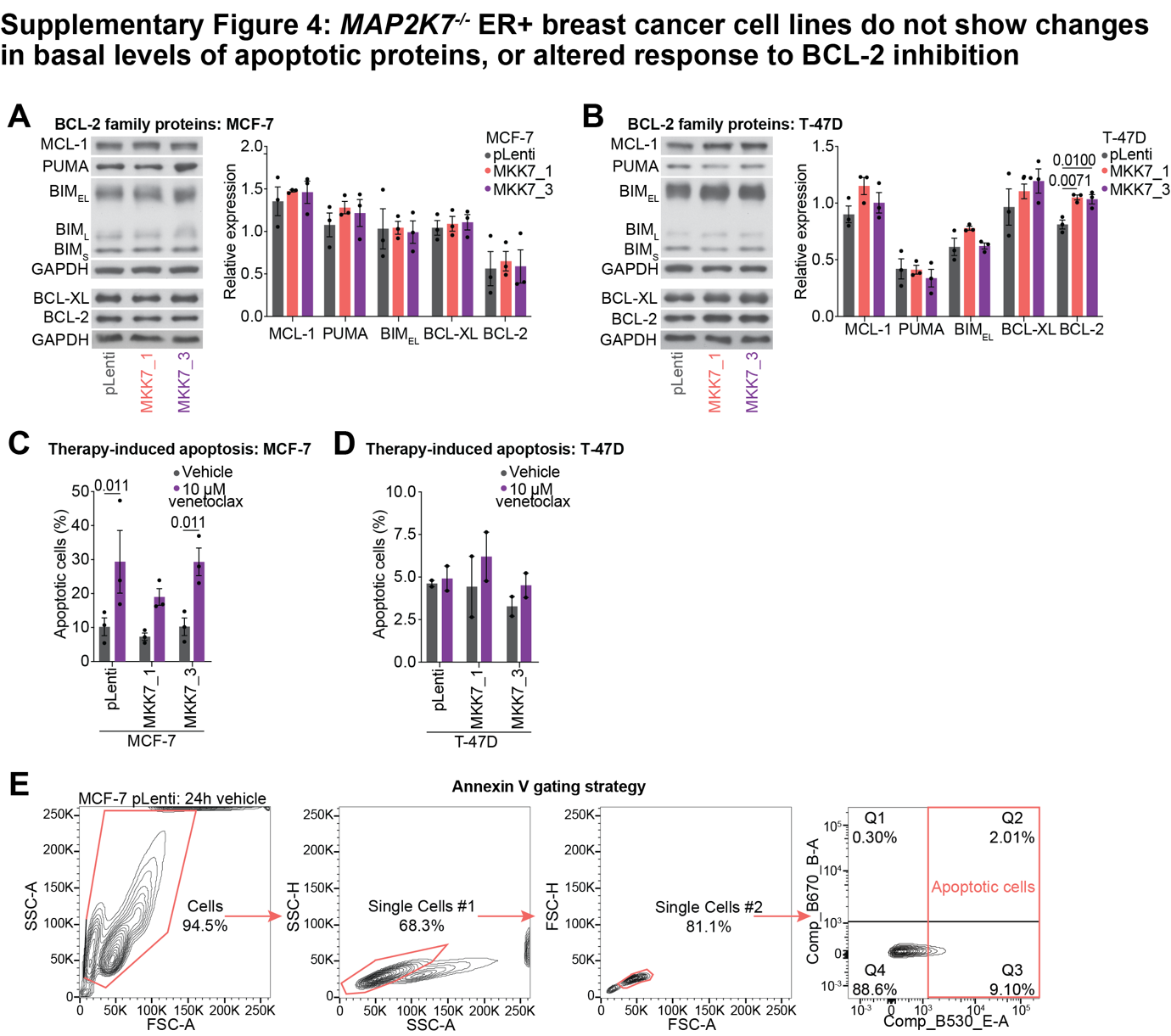

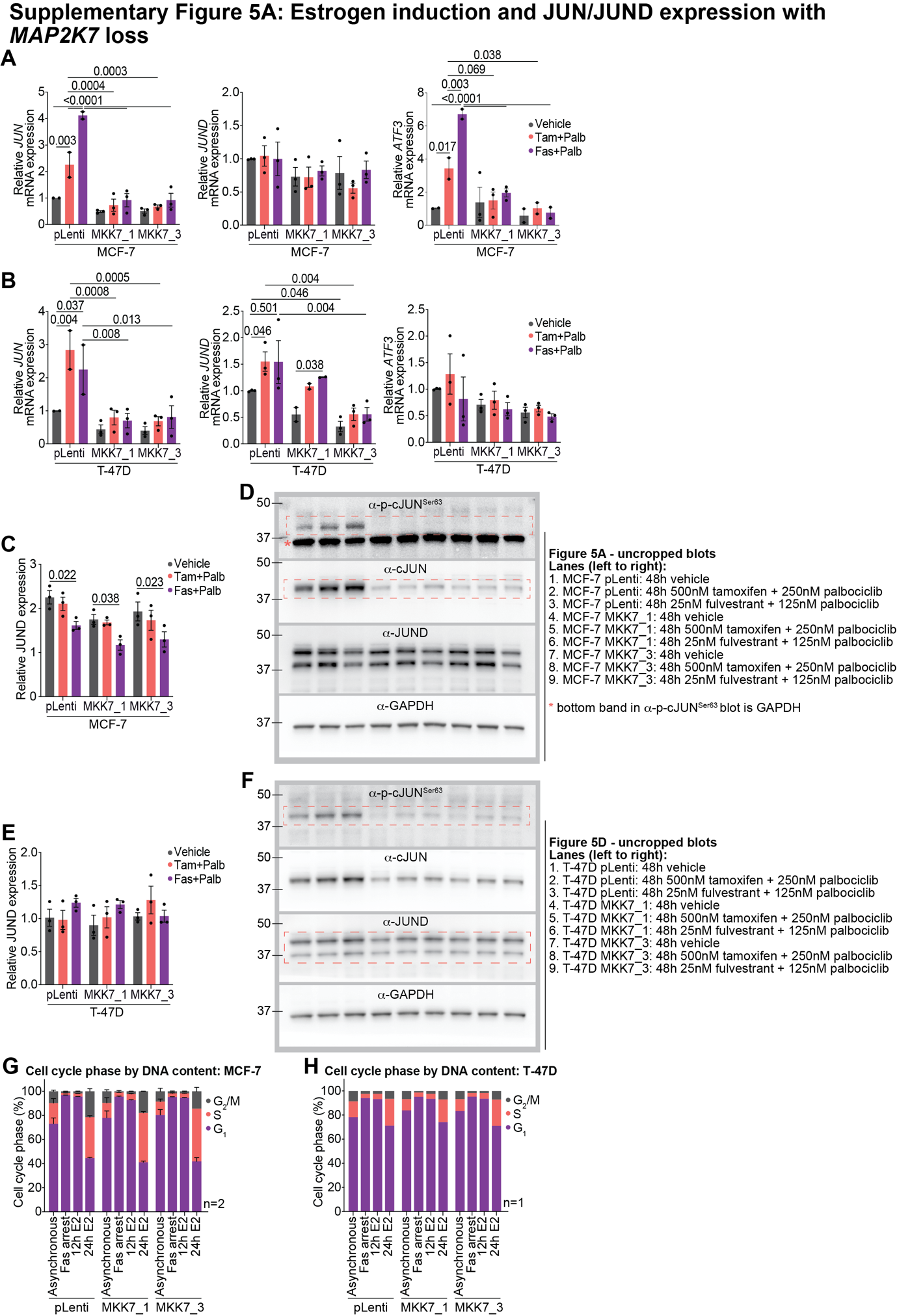

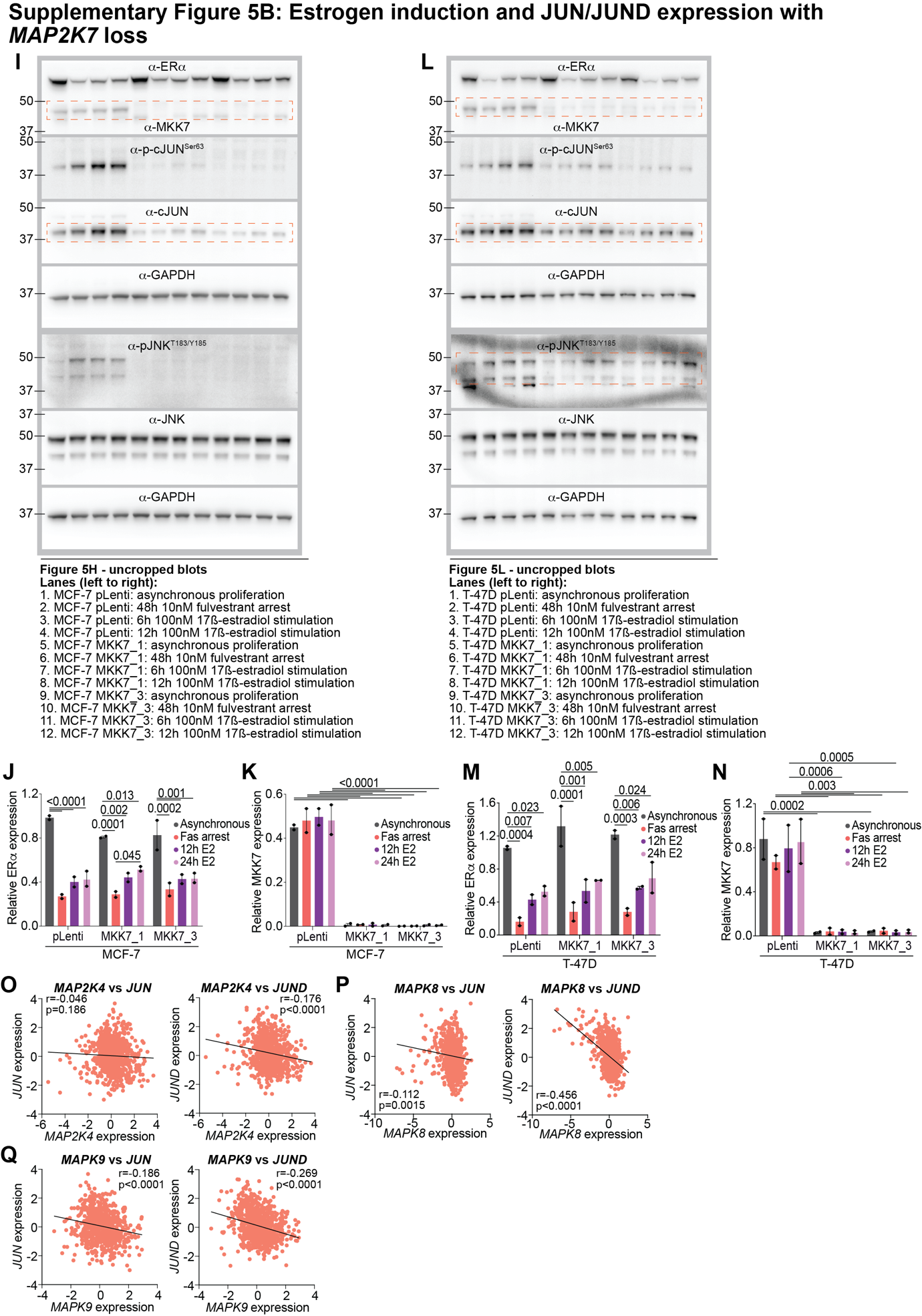

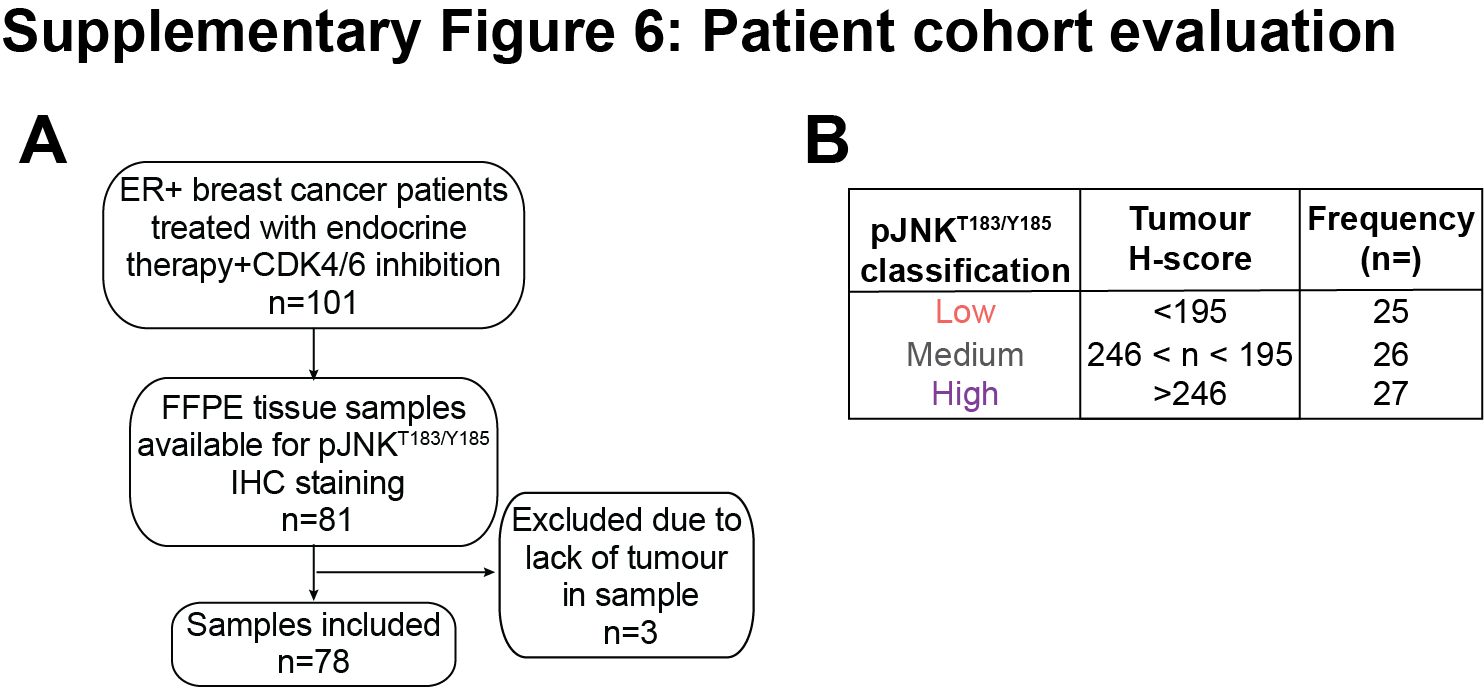
